# Generative design of programmable asymmetric β-barrel nanopores

**DOI:** 10.64898/2026.06.04.729630

**Authors:** Annika Philomin, Ria Sonigra, Sagardip Majumder, Hsien-Jui (Michelin) Lin, Yanjing Li, Fanglei Xue, Ryan D. Kibler, Claire K. Baldus, Eva Trapido, Amelie Medeiros, Brian Coventry, Asim K. Bera, Alex Kang, Johnny Mendoza, Manish Kumar, Yi Yang, David Baker

## Abstract

Protein nanopores are powerful tools for molecular sensing, sequencing, and separation, but designing pores with programmable function remains challenging. Native homo-oligomeric transmembrane β barrels (TMBs) are used for these applications, but their uniform lumens limit spatial resolution and analyte discrimination. Although monomeric TMBs can be designed using energy-based methods, these approaches remain highly manual and limited to structural design rather than function. Here, we present a generative AI framework for TMB design, with backbone and sequence design models trained on a curated distillation set. We characterized 48 designs spanning 0.7–1.5 nm in diameter, diverse lumen chemistries, and hydrophobic thicknesses. Crystal structures closely match the design models. We demonstrate that our method produces customizable nanopores for ion sensing, DNA translocation, and transport across polymer membranes.

## Introduction

Nanopore sensing has emerged as a transformative technology for single-molecule analysis, offering a direct, label-free window into the dynamics of DNA, RNA, and proteins. Monitoring the blockage of ionic current through a nanopore enables resolution of polynucleotide DNA sequences and conformational states of individual polypeptides(*1*, *2*). Nanopores also provide a route to molecular filtration, where precisely defined pore geometries and charges can govern size and chemistry-dependent separation of solutes(*3–6*). These efforts have largely relied on repurposing natural membrane proteins, including β-barrel pores such as α-hemolysin and MspA, as well as channel proteins such as aquaporins(*6–9*). However, their symmetric architectures create repetitive chemical environments within the pore lumen, limiting engineering of localized sensing zones and spatially defined interactions needed to improve sensing resolution and enable selective filtration of analytes(*8*, *10–12*). While monomeric bacterial porins like OmpF and OmpG offer a degree of structural asymmetry(*13*), they are burdened by evolutionary baggage, such as flexible gating loops which often lead to elevated noise levels and spontaneous gating(*13*). Their fixed pore architectures constrain reengineering without compromising pore integrity(*14–16*), limiting quantitative molecular sensing(*17*, *18*) and separation applications(*3*). *De novo* protein design offers a solution by enabling the construction of bespoke nanopores from first principles. Energy-based and parametric design strategies produced pore-forming TMBs with only a limited range of pore lining chemistries and geometries(*19–22*). Extending generative deep-learning methods, which have transformed soluble protein design, to TMBs has remained challenging because of the scarcity of TMB structures in the Protein Data Bank (PDB). Consequently, de novo TMB design has remained focused on structural design rather than on systematically programming pore properties for function.

We reasoned that current limitations in designing diverse *de novo* TMBs for nanopore applications could be addressed by an end-to-end generative AI pipeline capable of generating asymmetric barrels with tunable hydrophobic thickness, extracellular domains, and a broad range of pore constrictions (Fig. 1a). Here, we present a two-stage design pipeline for TMB design, consisting of separate backbone and sequence generation steps, using structure prediction tools to evaluate and filter designs *in silico*. Using this approach, we generated pores for ion sensing, DNA translocation, and ion transport across monoglyceride bilayer networks and synthetic block copolymer membranes(*3*) (Fig. 1b).

**Figure 1.**
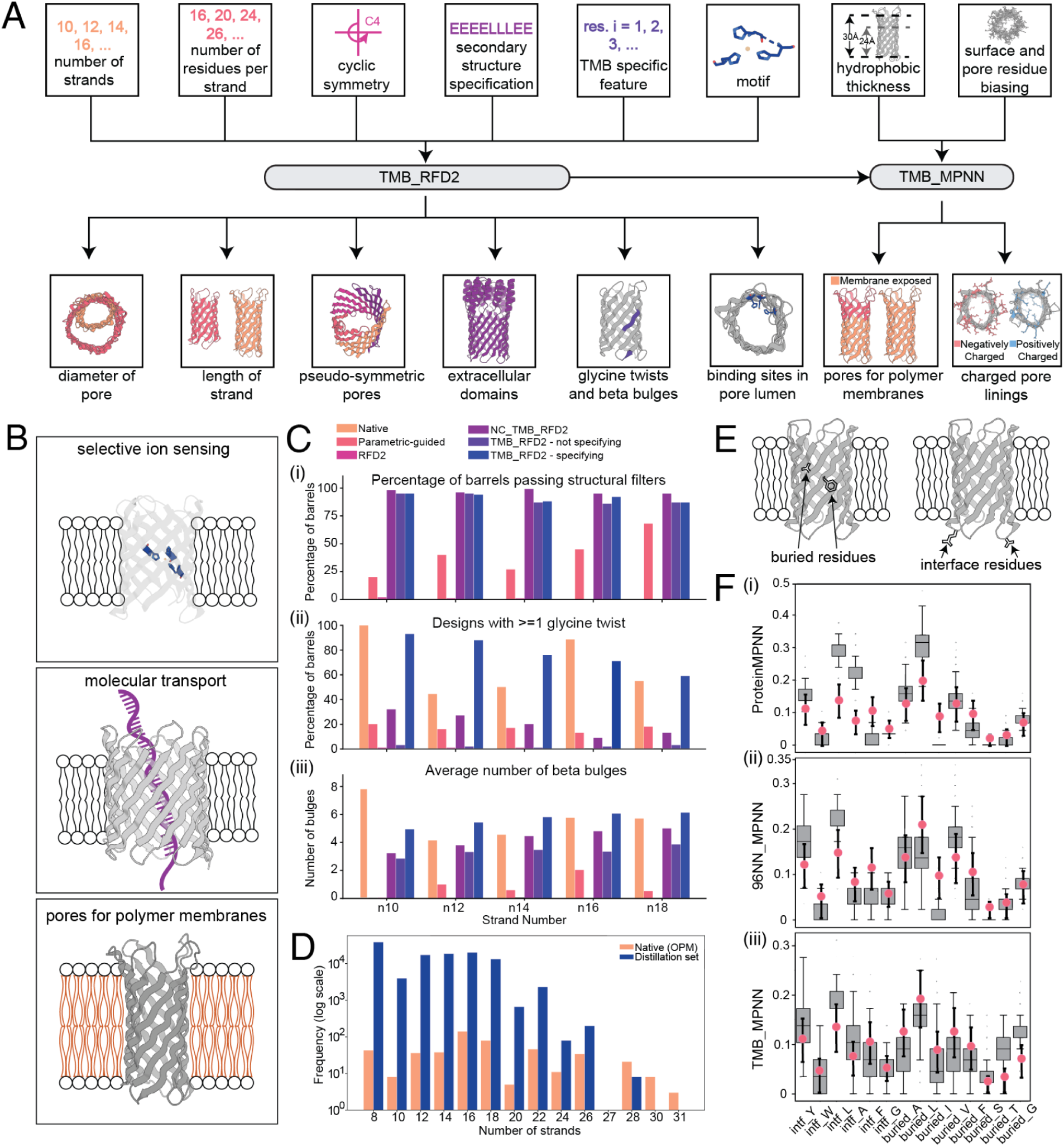
Overview of design space, applications, and new methods. **(A)** TMB_RFD2 generates diverse TMB geometries from inputs such as strand number, residues per strand, symmetry, secondary structure specification, glycine-twist strand placement, β-bulge placement, and motif residues. The resulting backbones are then passed through TMB_MPNN to generate sequences with customizable hydrophobic thickness and pore-lining chemistries. **(B)** Representative functional capabilities of deep learning *de novo* designed TMBs, including ion sensing, molecular transport, and pores compatible with polymer membranes. **(C)** Comparison of earlier methods with NC_TMB_RFD2 and TMB_RFD2. (i) Percentage of generated backbones passing β-barrel geometric filters across 100 backbones per strand count for parametric-guided diffusion, RFD2, NC_TMB_RFD2, and TMB_RFD2 with and without glycine twist and β-bulge constraints applied during inference. Native β-barrels are omitted because this metric evaluates the success rate of computational backbone generation and is therefore not directly applicable to experimentally determined native structures. (ii) Percentage of designs with at least 1 glycine twist in backbones generated by each method, including TMB_RFD2 designs generated with specified glycine-twist positions during inference. (iii) Number of β-bulges observed in backbones generated by each method, including TMB_RFD2 designs generated with specified β-bulge constraints during inference. **(D)** Comparison of the size of our curated distillation dataset with experimentally solved transmembrane β-barrel structures in the Orientations of Proteins in Membranes (OPM) database(*27*). Counts are shown on a logarithmic scale. **(E)** Representation of buried and interface residues **(F)** Surface amino-acid composition of sequences designed on robust energy-based β-barrel backbones(*19*, *21*) using (i) ProteinMPNN, (ii) 96NN_MPNN, and (iii) TMB_MPNN. Pink dots indicate amino-acid frequencies observed in native TMBs.

## Results

### Distillation set construction

Functional TMB design requires control over chemical and structural features. RFDiffusion2(*23*) (RFD2) can scaffold atom-level sites and handle ligands while retaining most features of RFDiffusion(*24*), making it an attractive starting point for designing ligand-sensing channels and customized pore constrictions for sequencing and other nanopore applications. However, when we tried generating TMBs with RFD2, control over barrel geometry was limited, barrel closure was poor, and the overall success rate in creating designable structures was low (Fig. 1c(i)). To improve TMB backbone generation, we curated a large TMB distillation set from the ‘’Is It A Barrel’’ database, using sequences shorter than 600 amino acids (*25*). Structures were obtained from the AlphaFold Database (AFDB) when available or predicted with AlphaFold2 (AF2), and then filtered by predicted local distance difference test (pLDDT) (>75), secondary-structure assignment, backbone hydrogen bonding, and geometric features (see Methods) (*26*). This reduced 485,000 predicted structures to 117,447 high-confidence monomeric TMBs, generating a dataset orders of magnitude larger than the 699 experimentally determined TMB structures in the Orientations of Proteins in Membranes (OPM) database(*27*) (Fig. 1d) and providing the scale needed for deep-learning-based TMB design.

### TMB_RFD2 fine-tuning and conditioning

We next finetuned RFD2 for TMB generation using the TMB distillation set. We utilized RFD2’s 1D residue-level secondary-structure feature that encodes α-helices, β-strands, and loops to capture the coarse-grained organization required for β-sheet formation. Building on the 2D residue-residue adjacency matrix used to encode spatial relationships, we tokenized β-strand pairing within this matrix to provide the model with explicit information about strand-strand interactions and global β-barrel topology. During training, we used selective masking to provide this information only for the middle beta strand residues, to prevent leakage of strand-length information, encouraging the model to infer β-sheet organization from context rather than explicit segment boundaries. This model is termed Non-Conditioned_TMB_RFD2 (NC_TMB_RFD2). It led to a substantial increase in the number of designs that satisfied the same β-barrel structure-based geometrical filters used to curate the distillation set (Fig. 1c(i)).

Since native transmembrane β-barrels use glycine twists(*19*, *28*) and β-bulges(*19*, *29*) to relieve strain arising from β-barrel closure and insertion into lipid membranes(*19*), we evaluated backbone quality based on the extent to which these features were incorporated into generated designs (Fig. 1c(ii, iii)). Glycine twists are local deviations in β-strand geometry caused by glycine residues which introduce bends that allow otherwise flat β-sheets to close into cylindrical β-barrels(*19*, *28*). β-bulges are localized irregularities near loops connecting antiparallel β-strands, where the hydrogen-bonding register skips a position and creates a small offset in the sheet that shapes barrel curvature(*19*, *29*). Analysis of structures generated using the NC_TMB_RFD2 model showed that they contained fewer glycine twists and β-bulges than structures in the PDB (Fig. 1c(ii, iii)) and our distillation set (Fig. S1).

To promote formation of these features we introduced 1D binary input features indicating the presence of glycine twists and β-bulges, and randomly masked these features during training to improve generalization. This model, termed TMB_RFD2, gives better quality backbones compared to NC_TMB_RFD2, with most designs having the desirable glycine twists and β-bulges (Fig. 1c(ii, iii)). These results show that conditioning on glycine twists and β-bulges increases incorporation of these native structural features in generated backbones.

Comparing TM-scores of backbones generated with TMB_RFD2 to those from previous methods(*22*) reveals substantially greater structural diversity in TMB_RFD2 outputs as is seen from lower pairwise TM-scores between designs across different strand numbers (Fig. S3). Notably, this structural diversity is achieved from minimal inputs, requiring only the strand number, average number of residues per strand, and optional specification of glycine twists and β-bulges (Fig. S4). Thus, TMB_RFD2 enables broad exploration of TMB backbone geometries without requiring extensive user-defined constraints.

### TMB_MPNN fine-tuning and conditioning

Previous *de novo* TMB design approaches have relied on Rosetta energy-based sequence design protocols(*19*, *21*), which are geometry-dependent and require tuning many interdependent parameters and constraints to achieve membrane-appropriate packing and hydrogen bonding. As a result, older TMB design methods relied heavily on expert knowledge and trial-and-error. The deep-learning based sequence design model ProteinMPNN(*30*) generated sequences with good structure prediction metrics for asymmetric TMB backbones designed by NC_TMB_RFD2 and TMB_RFD2, as assessed by AlphaFold2 and AlphaFold3. However, experimental characterization revealed no expression in *E. coli* (Fig. S6). This can likely be attributed to an enrichment of aliphatic residues in the buried region (Fig. 1e) of ProteinMPNN(*30*) sequences. On backbones from previously validated energy-based pores(*19*, *21*), A, V, L, and I were overrepresented in this region (Figs. 1f(i), S7), highlighting imbalances in sequence composition. Therefore, we next set out to adapt ProteinMPNN to enable generation of TMB sequences that fold and assemble experimentally.

We first retrained ProteinMPNN on the curated TMB distillation set with a strong bias on the distillation set structures as compared to the PDB. Initial benchmarking showed that sequences from this model better matched native TMB sequence patterns, with significantly lower hydrophobic amino-acid frequencies in the buried region (Fig. S8a). However, AlphaFold2 predicted very few designs to form TMB structures with high pLDDT scores and low Cα root-mean-square deviation (RMSD) values (Fig. S8b). Since TMBs lack a densely packed central core, especially at larger strand numbers, we reasoned that the model could not capture enough of the global structural features that encode the TMB fold. To address this, we increased the MPNN receptive field from 32 to 96 nearest neighbors, allowing residues on one side of the barrel to better sense residues on the opposite side. We retrained ProteinMPNN with this 96-neighbor receptive field on PDB structures alone, creating a baseline model for comparison to models trained with additional TMB-specific data. We refer to this model as 96NN_MPNN. We then fine-tuned 96NN_MPNN on the curated TMB distillation set to adapt sequence generation to the amino-acid patterns characteristic of native TMBs. We refer to this fine-tuned model as TMB_MPNN. Sequences generated by both 96NN_MPNN and TMB_MPNN showed good AlphaFold2 and AlphaFold3 structure prediction metrics. Compared to 96NN_MPNN, which still overrepresented aliphatic residues in the buried barrel region (Fig. 1f(ii)), TMB_MPNN produced more balanced amino acid compositions on backbones from known working designs generated by previous energy-based methods(*19*, *21*) (Figs. 1f(iii), S9). Together, these results support the use of TMB_MPNN for TMB sequence design, as it maintains strong structure prediction metrics while better recapitulating native TMB amino-acid composition.

### Design and experimental validation of TMBs

To determine which combination of backbone and sequence design models performed best in vitro, we first generated backbones using NC_TMB_RFD2 and TMB_RFD2 by specifying (i) strand number (10, 12, 14, or 16) and (ii) strand length (14–16 residues per strand). For TMB_RFD2 designs, glycine-twist placements were additionally specified on two to four strands per barrel. For a subset of TMB_RFD2 designs, β-bulge constraints were also specified during inference, although most designs were generated without explicit β-bulge constraints because the model frequently generated β-bulges autonomously (Fig. 1c(iii)). We next designed sequences on these backbones using 96NN_MPNN and TMB_MPNN. All candidate sequences generated were filtered using AlphaFold3 or RosettaFold3 as structure prediction oracles. In silico validation used stringent filtering criteria such that only designs exhibiting high pLDDT and minimal Cα RMSD from the target scaffold were prioritized for subsequent experimental characterization.

Proteins were expressed in *E. coli* as inclusion bodies, purified through detergent wash cycles, denatured, and subsequently refolded in detergent prior to characterization (see Methods). Sequences generated by 96NN_MPNN did not yield designs that expressed in *E. coli* (Fig. S12), despite high RosettaFold3 or AlphaFold3 confidence. Sequences designed by TMB_MPNN on NC_TMB_RFD2 backbones showed expression rates in *E. coli* ranging between 20-40% across different strand-number design campaigns (Fig. S13). In comparison, sequences designed by TMB_MPNN on TMB_RFD2 backbones expressed more robustly, with expression rates ranging from 60-80% across corresponding campaigns (Fig. S14). These data show that conditioning on native TMB features enables TMB_RFD2 to achieve higher expression success rates than NC_TMB_RFD2. Additionally, they show that TMB_MPNN represents a clear advance over both ProteinMPNN and 96NN_MPNN in expression.

To determine whether the generated TMBs were well folded, we performed size-exclusion chromatography (SEC). Across NC_TMB_RFD2 and TMB_RFD2 backbones with sequences generated by TMB_MPNN, a total of 42 10-stranded, 47 12-stranded, 91 14-stranded, and 101 16-stranded designs were tested (Table 1). SEC results showed that several designs generated with NC_TMB_RFD2 exhibited peaks at the expected elution volume (14-16 mL), consistent with well-folded proteins in detergent micelles. Although, many of these traces also showed prominent aggregate peaks (Fig. S17; Table 1). In comparison, designs generated with TMB_RFD2 showed a higher fraction of well-defined SEC peaks across all strand types, eluting at the expected monomeric position and exhibiting fewer dominant aggregation peaks (Fig. S18; Table 1). Although n18 designs generated with TMB_RFD2 were also tested, they did not produce well-defined SEC profiles (Fig. S21). Together, these results show that TMB_RFD2 generates better-folded designs than NC_TMB_RFD2 across 10-, 12-, 14-, and 16-stranded barrels.

**Table 1.**
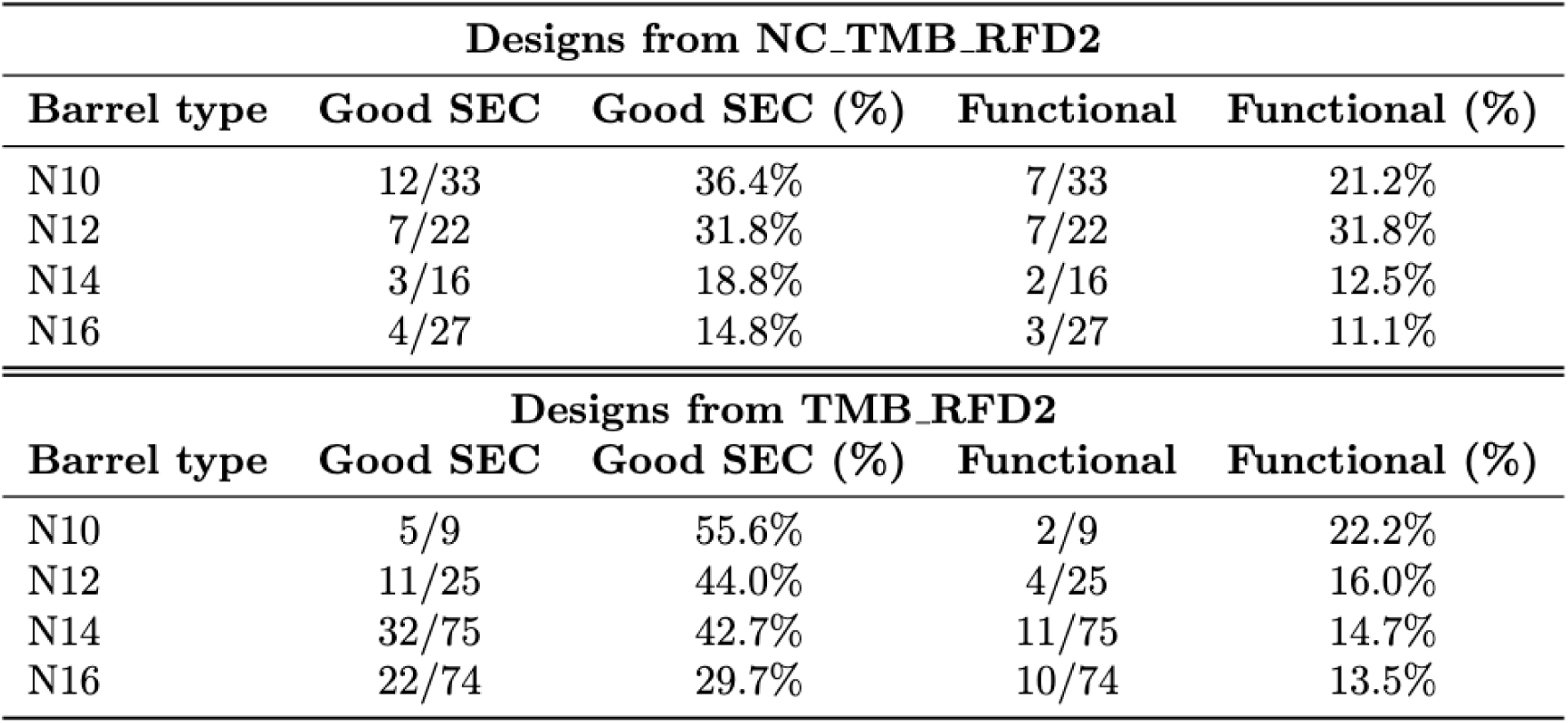
SEC quality and functional success rates for designed TMBs. Two design sets generated with different diffusion models, NC_TMB_RFD2 and TMB_RFD2, were compared after sequence design with TMB_MPNN. For each barrel type, the number of designs with good SEC profiles and functional success rates are shown relative to the total number of designs ordered.

| Designs from NC_TMB_RFD2 |  |  |  |  |
| --- | --- | --- | --- | --- |
| Barrel type | Good SEC | Good SEC (%) | Functional | Functional (%) |
| N10 | 12/33 | 36.4% | 7/33 | 21.2% |
| N12 | 7/22 | 31.8% | 7/22 | 31.8% |
| N14 | 3/16 | 18.8% | 2/16 | 12.5% |
| N16 | 4/27 | 14.8% | 3/27 | 11.1% |

**Table 1. SEC quality and functional success rates for designed TMBs.**
| Designs from TMB_RFD2 |  |  |  |  |
| --- | --- | --- | --- | --- |
| Barrel type | Good SEC | Good SEC (%) | Functional | Functional (%) |
| N10 | 5/9 | 55.6% | 2/9 | 22.2% |
| N12 | 11/25 | 44.0% | 4/25 | 16.0% |
| N14 | 32/75 | 42.7% | 11/75 | 14.7% |
| N16 | 22/74 | 29.7% | 10/74 | 13.5% |

We used HOLE(*31*) to analyze the lumen size and shape of several designs spanning n10–n16 barrels and found a diverse range of lumen architectures, with pore diameters ranging from 0.7-1.5 nm and varying constriction geometries (Fig. 2a). To assess whether the designed TMBs conduct ions, we performed single-molecule electrophysiology experiments using the Orbit 16 TC platform (Fig. 2a). Upon addition of purified proteins to planar 16-carbon DPhPC lipid bilayers, we observed stepwise increases in ionic current, where each discrete jump corresponds to the insertion of a single pore into the membrane (Fig. 2a). Histograms derived from single-channel recordings revealed stable, uniform open-state conductances with low stochastic noise under both positive and negative voltage bias (Fig. 2a). Several designs showed different current distributions at positive and negative voltages, suggesting asymmetric transport behavior within the pore (Fig. 2a).

**Figure 2.**
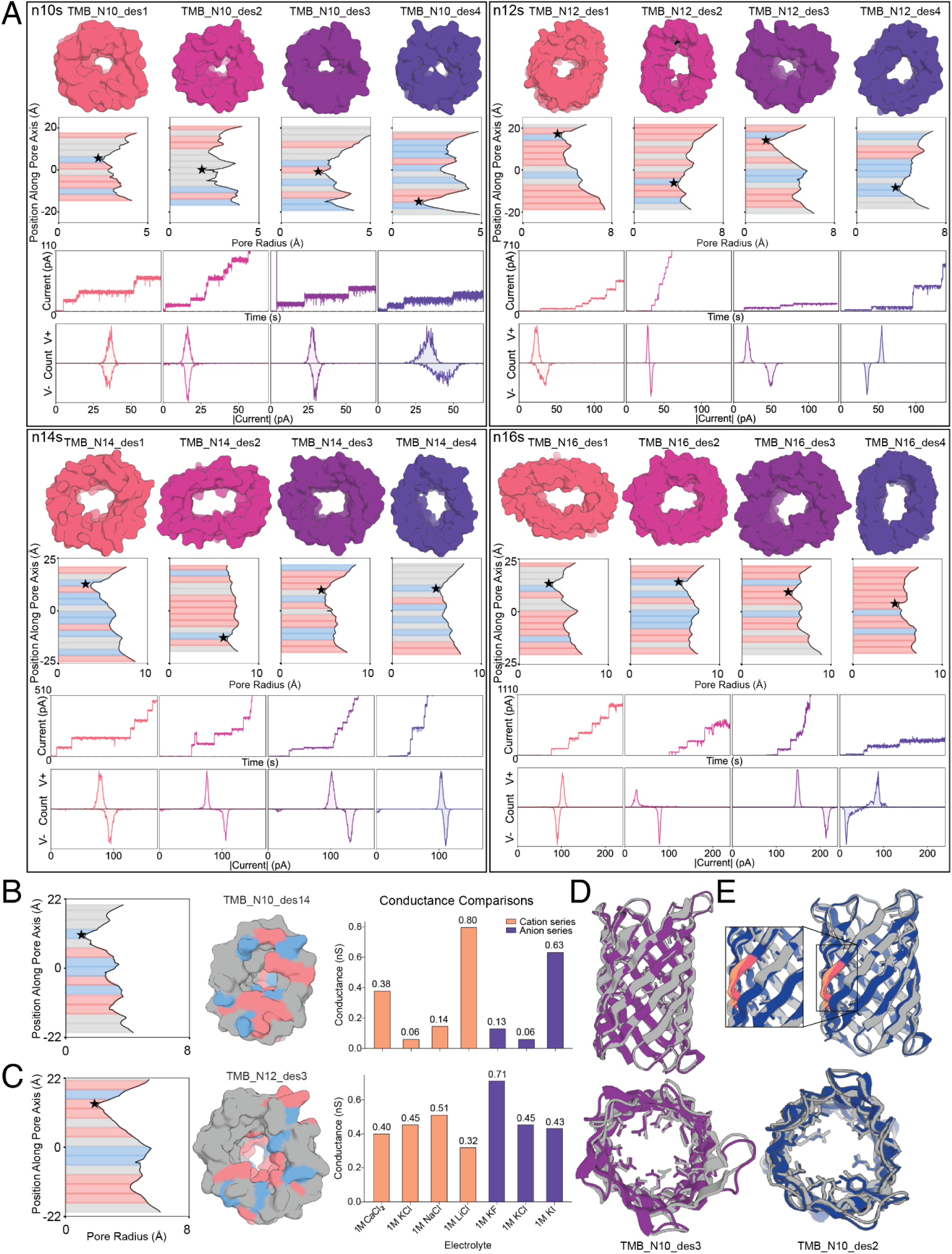
Experimental validation of *de novo* TMB designs. **(A)** Electrophysiological characterization of designed 10-, 12-, 14-, and 16-stranded β-barrels in planar lipid membranes. Top views of representative structures are shown together with the net charge of the pore lining (pastel pink, negative; pastel blue, positive) and constriction sites (black stars). Representative insertion (jump) events were recorded at applied potentials of 50 or 100 mV (red lines) in lipid bilayer membranes in 1M or 1.5M KCl. Conductance histograms were compiled from single-channel recordings at ±50 or ±100 mV. **(B-C)** Ion transport measurements for **(B)** a 10-stranded pore (net charge -1.8) and **(C)** a 12-stranded pore (net charge -3.8). Left: radius profiles and top views of the pores. Right: conductance comparison of the two pores measured in different cation and anion solutions. **(D)** Crystal structure (gray) of TMB_N10_des3 (Cα RMSD = 1.15 Å) without specified glycine twists aligned with the design model (purple). Side and top views are shown, respectively, with selected pore-facing side chains displayed that are in close agreement between the crystal structure and the design model. **(E)** Crystal structure (gray) of TMB_N10_des2 with glycine twists specified on strands 2 and 8 (Cα RMSD = 0.726 Å) aligned with the design model (blue). Glycine twist residues are highlighted in orange in the crystal structure and pink in the design model. A zoomed-in view highlights the glycine twist on strand 2.

Across 10-, 12-, 14-, and 16-stranded barrels, NC_TMB_RFD2 and TMB_RFD2 produced comparable percentages of functional pores, defined by measurable conductance in planar lipid bilayers. NC_TMB_RFD2 designs ranged from 11.1-31.8% functional, while TMB_RFD2 designs ranged from 13.5-22.2% functional (Table 1). However, conductance alone does not reflect pore quality. Larger NC_TMB_RFD2 designs, particularly n14 and n16 barrels, often showed noisy pore-current behavior (Fig. S22). In contrast, TMB_RFD2 produced more robust n14 and n16 pores with reduced current noise (Fig. 2a, S24). These results suggest that TMB_RFD2 improves functional behavior in larger barrels, while also indicating that further stabilization may be needed for larger barrel architectures.

We therefore investigated whether differences in native-like features could help explain the improved performance of TMB_RFD2 designs. Proline residues are enriched at cis β-bulges in native TMBs (Fig. S23a) and can promote strand turning by disrupting the β-strand hydrogen-bonding pattern through the absence of a backbone NH(*32*). TMB_RFD2 backbones showed a higher fraction of TMB_MPNN-designed sequences containing prolines at cis β-bulges compared to NC_TMB_RFD2 backbones (Fig. S23b). To test whether this difference could contribute to pore performance, we introduced proline mutations at cis β-bulges in poorly functional NC_TMB_RFD2 designs. Indeed, these point mutations converted multiple previously non-functional NC_TMB_RFD2 designs into conductive pores (Fig. S23c). This data suggests that proline incorporation at cis β-bulges improves folding and pore functionality. Therefore, the higher frequency with which TMB_MPNN places prolines at cis β-bulges in TMB_RFD2 designs likely contributes to their improved performance.

To determine whether designed pores exhibited different ion-transport properties, we further characterised designs by measuring their ionic conductance with different cations and anions for one 10-stranded (Fig. 2b) and one 12-stranded (Fig. 2c) β-barrel pore with net charges of -1.8 and -3.8, respectively (Fig. S26). Ion conductance in these pores does not follow the usual bulk-electrolyte mobility trends based on hydration radii(*33–35*) (K⁺ > Na⁺ > Li⁺ ∼ Ca^2+^ for cations and I⁻ ∼ Cl⁻ > F⁻ for anions). Ion transport through nanopores often deviates from these trends because permeation depends not only on bulk mobility but also on electric field, pore electrostatics, hydration shell energetics, steric confinement, and specific ion-pore interactions(*36–41*). The low conductance in CaCl₂ for both pores is consistent with the strong hydration and high dehydration cost of Ca²⁺, which hinder transport despite its larger charge. The 10-stranded pore’s constriction with a diameter of ∼4 Å, is close to the hydration radius of Li^+^ ion (∼ 3.8Å), potentially explaining the preference for partially hydrated Li^+^ ions transport(*42*). The wider and more negatively charged 12-stranded pore favours the more mobile Na⁺ ion. In the anion series, the 12-stranded pore shows highest conductance in KF, suggesting that in this pore F⁻ is better accommodated, while larger I⁻ is slowed by steric or interaction-based effects. Overall, these results demonstrate that changes in β-barrel architecture and lumen charge distribution can substantially reshape ion transport behaviour.

The structural integrity of nine designs was further corroborated by circular dichroism (CD) measurements (Fig. S27), which confirmed β-sheet secondary structure with a characteristic minimum near 215 nm. Two 10-stranded designs were further validated by X-ray crystallography (Fig. 2d, e), providing atomistic confirmation of the designed topologies. One structure originated from a NC_TMB_RFD2 backbone design containing three cis β-bulges and no glycine twists. The crystal structure closely matched the 160-amino-acid backbone generated by NC_TMB_RFD2 (Cα RMSD = 1.15 Å) (Fig. 2d). It recapitulates the round pore shape observed in the top view due to the lack of glycine twists. The second structure was from a TMB_RFD2 backbone in which glycine twists were specified on strands 2 and 8. Similarly, the crystal structure closely matched the designed 160-amino-acid backbone (Cα RMSD = 0.726 Å), recapitulating the specified glycine twists on strands 2 and 8, consistent with the constraints imposed during inference (Fig. 2e). These glycine twists give rise to a more oval pore shape, which is likewise captured in the crystal structure. The design also has 4 cis β-bulges, in agreement with the crystal structure. These structures validate the ability of NC_TMB_RFD2 and TMB_RFD2 to generate experimentally accurate β-barrel topologies, including features like cis β-bulges and glycine-twists.

### Ion sensing via motif-scaffolding within TMBs

We explored the possibility of generating ligand-gated TMBs by introducing binding motifs into the lumen of our *de novo* designed TMBs. To enable the scaffolding of functional motifs for ion binding and active-site incorporation, we adapted the RFD2(*23*) motif scaffolding training protocol to randomly sample strand segments facing the pore interior during training. We then generated 10-stranded TMBs containing a three-histidine (His₃) copper binding motif (Fig. 3a). We selected Cu(II) as a test ligand because copper has well-characterized protein-binding motifs, making it a convenient model for testing whether metal binding can modulate ionic current and enable ligand-responsive pore design. We selected the His₃ motif because it creates a compact coordination environment compatible with the steric limits of the pore lumen^17–20^.

**Figure 3.**
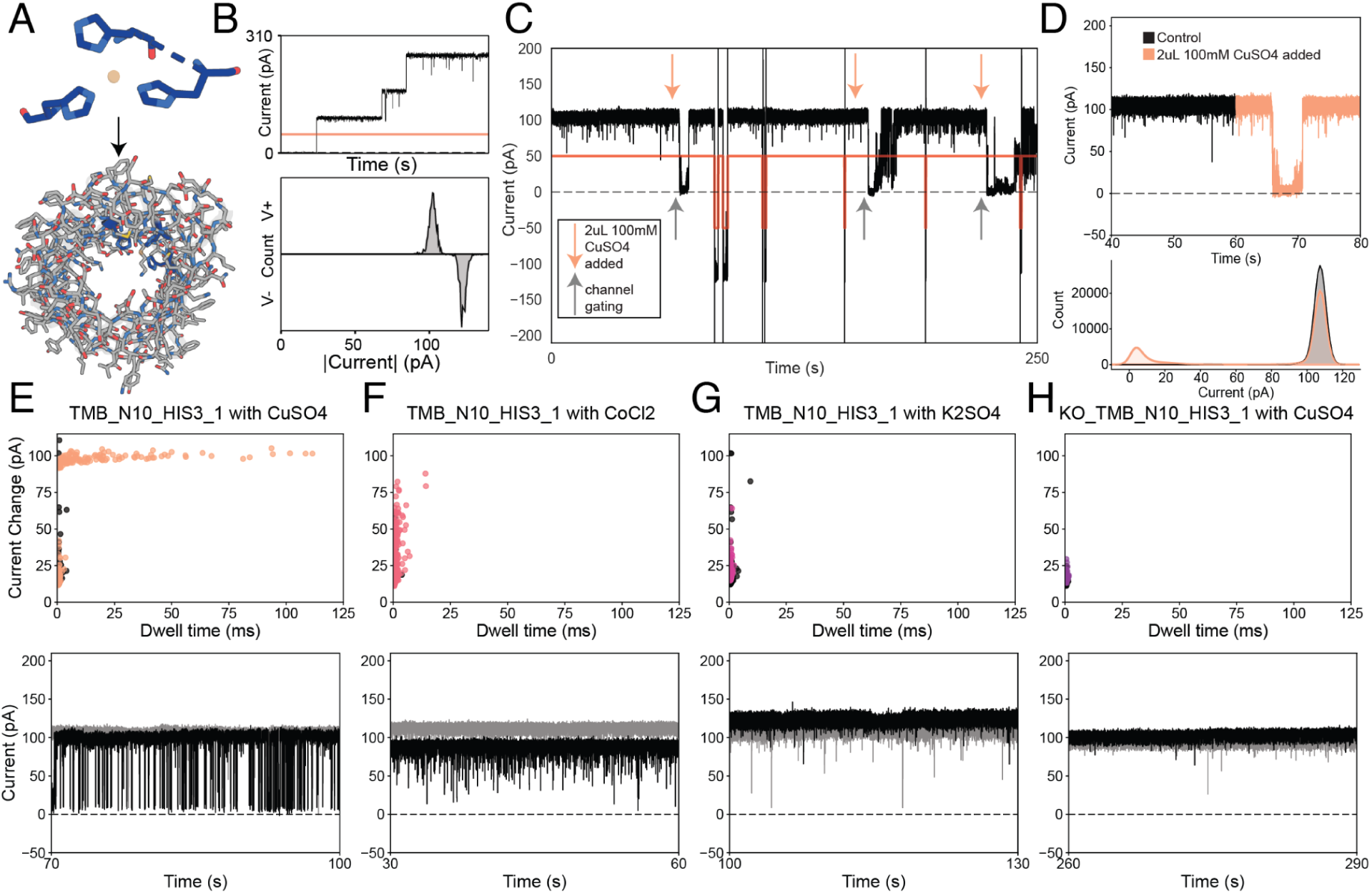
Ion sensing via motif-scaffolding within TMBs. **(A)** Schematic representation of the motif-scaffolded copper-binding site positioned within the pore lumen. **(B)** Insertion events of TMB_N10_HIS3_1 at 50mV applied (red line), and single-channel current histogram under 50mV and -50mV applied. **(C)** Representative current trace of TMB_N10_HIS3_1 showing three sequential additions of 2 µL of 100 mM CuSO₄ to a 200 µL recording chamber (orange arrows), followed by channel gating events (gray arrow). **(D)** Top: zoom-in on the 40-80 s window from panel C, before and after addition of 2 µL of 100 mM CuSO₄. Bottom: current histogram from the zoomed-in trace. **(E-H)** Scatter plots of dwell time (ms) versus current change (pA) relative to the open-pore current under control conditions and after ion addition, shown alongside representative raw current traces. Panels E-G show TMB_N10_HIS3_1, while panel H shows the knockout design, KO_TMB_N10_HIS3_1. Control conditions are shown as light gray traces and gray dots, while ion-incubated conditions are shown as black traces with colored dots.

Experimental characterization of a representative His₃ motif design, TMB_N10_HIS3_1, exhibited consistent insertion events and a well-defined open-state conductance in single-channel recordings in 1 M KCl (Fig. 3b). It further revealed copper-dependent gating immediately after addition of 2 µL of 100 mM CuSO₄ to the 200 µL chamber (Fig. 3c). Repeated additions produced reproducible and progressively amplified changes in channel activity (Fig. 3c). The shift in behavior is also evident when comparing histograms of current (pA) before and after CuSO₄ addition, which show a pronounced increase in events centered near 0 pA (Fig. 3d). After three sequential additions of 2 µL of 100 mM CuSO₄, resulting in a final concentration of 2.65 mM in the chamber, and following a 5-15 minute incubation period to allow diffusion and equilibration, gating became markedly more pronounced (Fig. 3e). Plotting dwell time versus current change relative to the open-pore current showed that CuSO₄ addition shifted events toward longer dwell times and larger closures compared with control recordings without CuSO₄ (Figs. 3e, S30). In contrast, addition of the same final concentration of CoCl₂ or K₂SO₄ in independent experiments did not produce comparable gating behavior, with dwell times and current changes remaining similar to those observed in the control recordings (Figs. 3f, g; S30). These results support the metal specificity of the scaffolded His₃ coordination site.

To further test whether the histidine motif was responsible for CuSO₄-induced gating, we generated a knockout variant of TMB_N10_HIS3_1, termed KO_TMB_N10_HIS3_1, by mutating one histidine to serine and another to methionine within the His₃ motif. Plotting dwell time versus current change relative to the open-pore current showed that KO_TMB_N10_HIS3_1 did not exhibit comparable long-lived closures after CuSO₄ addition (Fig. 3h), supporting that the scaffolded histidines are required for the observed gating behavior. These results demonstrate that molecular sensing can be encoded directly into de novo designed nanopores through placement of binding motifs.

### Molecular transport and DNA interactions with designed TMBs

To test whether larger designed pores could interact with DNA analytes, we evaluated two n14 pores. Using TMB_N14_des12, generated with TMB_RFD2 and TMB_MPNN, we measured ionic currents before and after addition of a 22 repeat thymine base ssDNA analyte (Fig. 4a). Recordings exhibited deep current blockades, whose duration decreased with increase in applied voltage indicating possible translocation through the pore. However, the amplitude of the blockades were very close to the open-state current for this pore, making it unlikely that sufficient resolution could be obtained for nanopore-based DNA sequencing(*10*). Another design, TMB_N14_des1, also generated with TMB_RFD2 and TMB_MPNN, had a larger effective constriction radius and pore-volume and was reasonably quiet at high voltages (Fig. 4b). This design had blockage amplitudes reaching 70-80% of the open-pore current in the presence of the same ssDNA analyte and a motor-enzyme (Fig. 4c), suggesting that translocation of DNA with a motor-enzyme could potentially generate sufficient variation in the signal to achieve resolutions needed for efficient base-calling. Towards this end, we investigated enzyme-driven transport by adding a premixed adapter-ligated sample, containing an in-house PET29b-based plasmid vector and sequencing buffer from a MinION rapid sequencing kit, on top of the pore. Clear DNA capture was observed and variations in current blockades showed continuous enzymatic activity and likely DNA threading (Fig. 4c). The *de novo* pore signal (Fig. 4d), compared with the true sequencing signal recorded for the same plasmid using an R10.4 pore-containing flow cell(*47*, *48*) (Fig. 4e), exhibited noisier conductance states and more uneven dwell times at each step. However, the maximum current blockade was very close to the open-state current, indicating a smaller constriction opening compared to the R10.4 pore. State-transition probabilities for TMB_N14_des1, obtained by fitting a Hidden Markov model (HMM) to the low-pass-filtered signal (Fig. 4d), are consistent with enzyme-mediated state transitions, with nonzero stepping probabilities in both the forward and backward directions. To our knowledge, this is the first published current trace of enzymatic DNA threading through an asymmetric nanopore with no soluble domain. Corresponding state-transition probabilities for the true sequencing signal from the R10.4 pore, obtained using an equivalent HMM fit, are shown in Fig. 4e. Further optimization of our designs by modifying pore chemistry, varying constriction radii, and adding a custom helicase-stabilizing domain could increase single-molecule sensitivity beyond what is currently achievable with modified natural nanopores.

**Figure 4.**
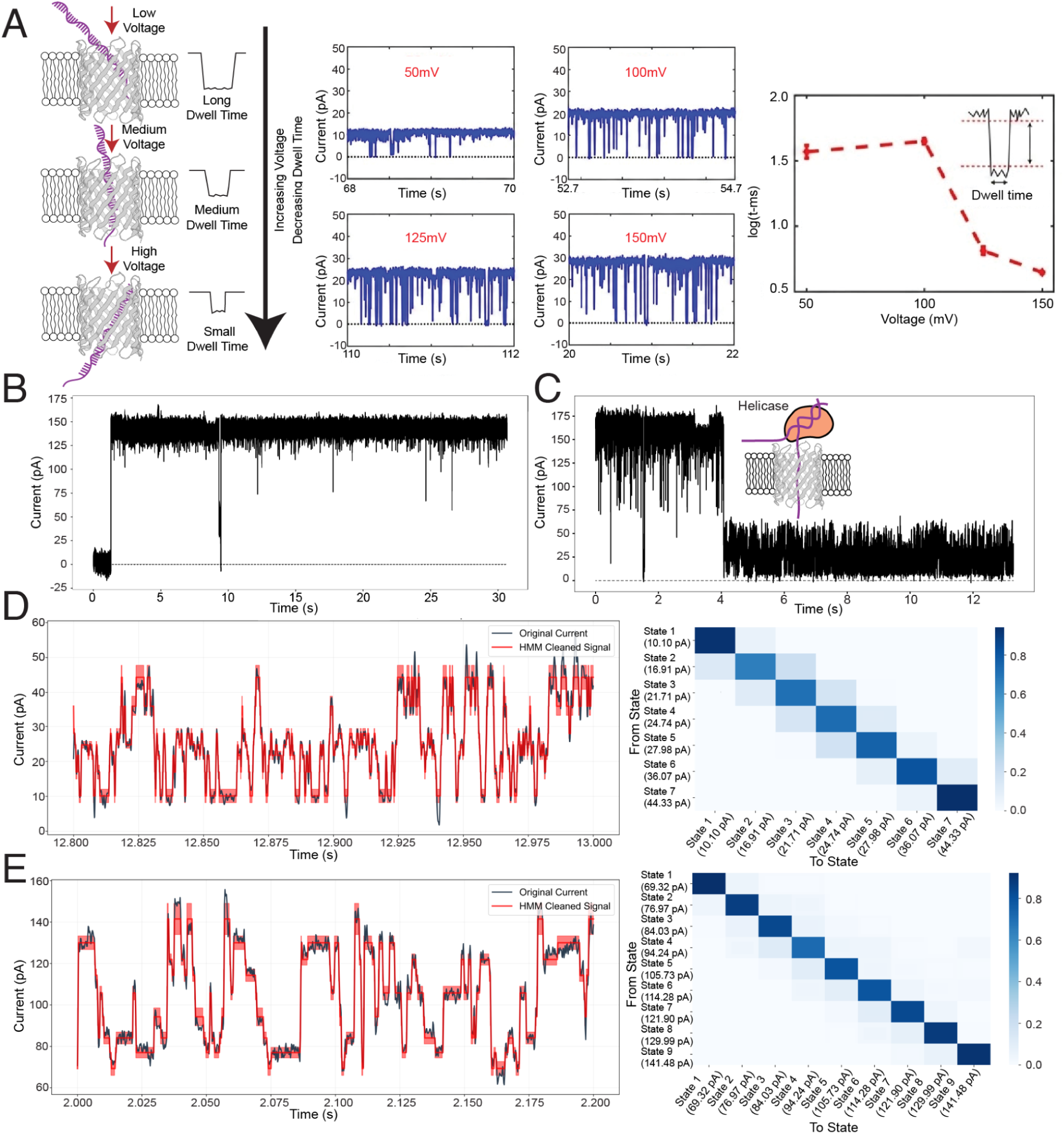
Molecular transport and DNA interactions with designed TMBs. **(A)** Schematic depicting the voltage dependence of dwell time for indirect measurement of charged analyte translocation. Current traces of TMB_N14_des12 with added ssDNA (22 Thymine repeats) show DNA-dependent blockages in 200mM NaCl and the plot of log (dwell time) vs voltage depict a translocation regime for the ssDNA past an applied voltage of 100mV.**(B)**Current trace of TMB_N14_des1 in 500mM KCl in the absence of DNA **(C)** Current trace of a captured helicase-loaded DNA strand. **(D)** Left: 250 ms current trace pertaining to roughly 100 bases filtered at 2 kHz showing enzyme mediated DNA threading across TMB_N14_des1. Right: State transitioning-probability heatmap of a trained hidden Markov model with 7 states as optimised using minimal AIC criterion**. (E)** Left: 250 ms current trace pertaining to roughly 100 bases filtered at 2kHz showing enzyme mediated DNA threading across a R10.4 MinIon flow cell pore. Right: State transitioning-probability heatmap of a trained hidden Markov model with 9 states as optimised using minimal AIC criterion.

### Design of TMBs for Polymer Membrane Integration

A major bottleneck in the commercial translation of nanopore technology is the mechanical fragility of lipid bilayers. Synthetic block copolymer membranes offer enhanced stability and are promising platforms for membrane separation applications(*3*), yet they are significantly thicker than biological bilayers (∼40 Å vs. ∼23 Å). This creates a thermodynamic barrier known as hydrophobic mismatch that natural proteins cannot easily overcome (Fig. 5a). Functional incorporation of native membrane proteins into polymer membranes is non-trivial and can require extensive empirical tuning of reconstitution conditions(*49*, *50*).

**Figure 5.**
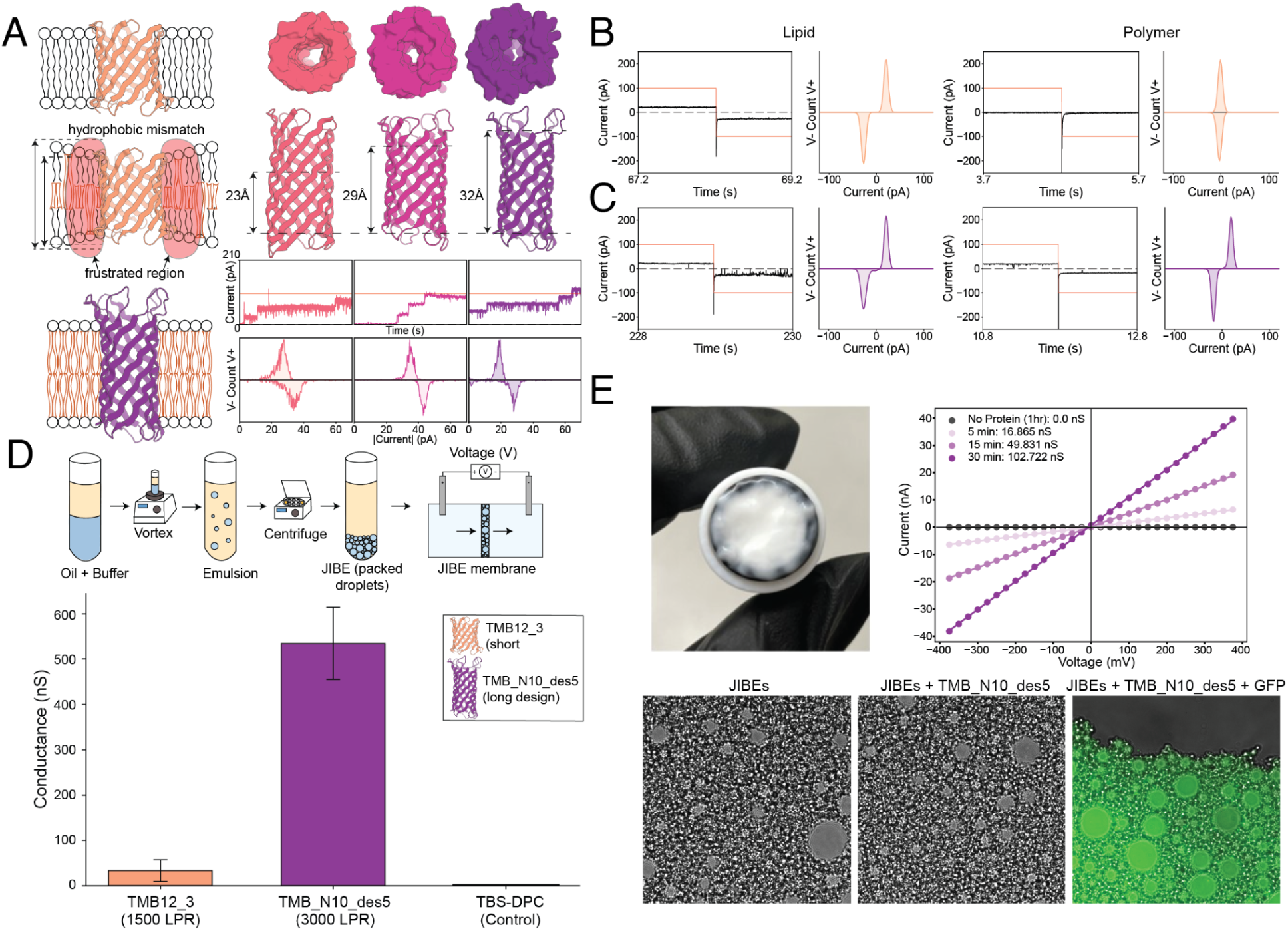
Tuning hydrophobic thickness of TMBs enables conductance in polymer membranes and macro-scale membrane mimetics. **(A)** Schematic showing hydrophobic mismatch between a design with short hydrophobic thickness and a polymer membrane, and hydrophobic matching for a design with longer hydrophobic thickness. Representative stepwise insertion events recorded in lipid bilayer membranes in 1.5M KCl at an applied potential of 100 mV (red lines) and corresponding single-channel current histograms recorded at ±100 mV for longer designs with varying hydrophobic thicknesses (black dashed lines). Two designs, shown in pink and purple, were conditioned with TMB_MPNN to have greater hydrophobic thicknesses than conventional lipid-inserting TMBs. **(B-C)** Comparison of **(B)** a short design, TMB12_3, from previous methods(*21*) to **(C)** a longer design with increased hydrophobic thickness, TMB_N10_des5, made with TMB_RFD2 and TMB_MPNN. Single-channel recordings at ±100 mV (red lines) in 1 M KCl showed that both designs insert into lipid membranes. In PBd-b-PEO polymer membranes, TMB_N10_des5 inserts, whereas the short-barrel design does not. **(D)** Schematic of the JIBE setup and comparison of bulk membrane conductance. Bulk conductance measurements in C18 monoglyceride bilayers show that TMB_N10_des5 exhibits substantially higher conductance than TMB12_3. **(E)** I-V traces show increased conductance over time after TMB_N10_des5 addition to JIBEs. Brightfield microscopy images show gently pressed JIBEs without and with TMB_N10_des5. Image of JIBEs with TMB_N10_des5 and GFP show a high density of lipid-coated droplets of varying sizes by fluorescence microscopy.

To generate barrels compatible with membranes composed of synthetic block copolymers, we trained a hydrophobic thickness-aware variant of TMB_MPNN. This model is conditioned on hydrophobic thickness by tokenizing buried membrane-spanning regions and interfacial regions separately, enabling hydrophobic residue patterning to be tuned for membranes with different thicknesses and compositions.

To design TMBs compatible with thicker, polymer membranes, we used TMB_RFD2 to sample ∼20 residues per strand, rather than the 14 or 16 residues used previously. We then designed sequences for these backbones using our hydrophobic thickness-aware variant of TMB_MPNN. We allowed the model to either freely generate sequences or explicitly specified a target hydrophobic thickness of ∼30 Å. Using this approach, we identified functional designs with hydrophobic thicknesses ranging from ∼23 Å to 32 Å. (Fig. 5a). These constructs spontaneously inserted into planar 16-carbon DPhPC lipid bilayers and exhibited single-channel conductance with a defined open state (Fig. 5a).

We next tested whether tuning hydrophobic thickness could enable designed TMBs to function in synthetic block copolymer membranes. We compared TMB12_3, a pore generated using previous energy-based methods(*21*) and designed for lipid membranes with a ∼23 Å hydrophobic thickness, with TMB_N10_des5, a longer design from our new methods designed to have a ∼32 Å hydrophobic span. Both designs inserted and formed functional channels in 16-carbon DPhPC lipid membranes (Fig. 5b, c). In contrast, in membranes composed of the PEO-b-PBO block copolymer, only TMB_N10_des5 inserted and produced a stable single-channel conductance trace (Fig. 5c), whereas TMB12_3 failed to insert (Fig. 5b). Insertion in polymer membranes represents an important capability that was not achieved with previous de novo TMB design approaches.

To further test whether longer designed TMBs can function in thicker membrane environments, we evaluated TMB_N10_des5 in the JIBE droplet bilayer tissue platform(*51*) (Fig. 5d). This platform forms three-dimensional networks of monoglyceride bilayer-enclosed aqueous droplets, allowing multiple pores to be reconstituted and ionic currents to be measured in bulk (see Methods). TMB_N10_des5 produced clear insertion events and sustained high-conductance states, whereas TMB12_3 showed substantially lower conductance despite being tested at a lower lipid-to-protein ratio (LPR) (Fig. 5d). Conductance increases over time in the presence of TMB_N10_des5 as pore-containing JIBEs establish conductive pathways (Fig. 5e). Brightfield and fluorescence images show no visible differences between JIBEs with and without pores (Fig. 5e), indicating that the increase does not arise from droplet disruption. An image of JIBEs with TMB_N10_des5 and GFP shows the droplet volume and packing (Fig. 5e).

These results show that longer β-barrel nanopores with increased hydrophobic thickness can function in polymer membranes and thicker monoglyceride bilayers, broadening the membrane environments and recording platforms compatible with designed nanopores. This extended pore length and hydrophobic thickness can enable longer interaction pathways for analytes and hence increased selectivity during transport(*52–54*), with considerable potential for downstream filtration and molecular separation applications.

## Discussion

We describe a generative deep-learning driven framework for the *de novo* design of TMBs that expands both the accessibility and functional scope of prior approaches, enabling rapid, day-scale generation of experimentally tractable pores. TMB_RFD2 further provides accessible geometric control by allowing glycine twists and β-bulges to be specified during design, enabling barrel curvature and pore morphology to be shaped as desired. Designs can be obtained using randomly sampled glycine-twist placements, reducing the need for extensive manual specification. Together, these advances generate designs spanning a broad range of strand numbers (n = 10–16) and constriction geometries, while maintaining robust refolding and membrane insertion.

The ability of our approach to directly scaffold motifs within the pore lumen, exemplified by the design of a Cu-gated channel, enables the design of programmable TMBs that host precise functional elements. Our larger designed pores can also perform functional behaviors such as DNA translocation, illustrating the stability and functional robustness of the designed β-barrels. By implementing hydrophobic thickness dependent sequence-design, we enable the design of pores for a wide range of membranes beyond the phospholipid bilayers in nature, such as polymer membranes. Together, our deep learning framework enables the simple and modular design of β-barrel nanopores, expanding our ability to build and engineer membrane-spanning protein architectures that have historically been difficult to access.

At the same time, our results highlight current limitations of *de novo* TMB design and suggest directions for improvement. Scaling to larger architectures (n ≥ 16) led to less well-defined SEC profiles (Figs. S18, S22) and more heterogeneous conductance jumps, potentially due to reduced hydrogen bonding in the lumen of the barrel. In an effort to stabilize larger barrel designs, we used TMB_RFD2 to generate symmetry-enforced single-chain TMBs with short α-helical extracellular domains. Although these designs formed pores spontaneously, their noisy conductance traces suggest that the soluble helical domains were not sufficiently stabilized (Fig. S30). Future designs may require longer and more extensively packed α-helical domains, with fewer flexible or unstructured regions. For regular TMB pores, some designs showed stochastic gating despite the absence of natural TMB-like loop domains (Fig. S32). This suggests that future design efforts may need to more explicitly account for membrane context and barrel dynamics, for example through molecular dynamics-informed constraints or stability objectives during generative sampling(*55*).

Looking forward, the ability to rapidly generate asymmetric, functional TMBs places several previously aspirational applications within immediate reach. The demonstrated robustness of TMBs with longer hydrophobic thicknesses in polymer-lipid mixtures, combined with their portability across electrophysiology platforms, positions these designs as attractive components for industrial membrane integration and filtration systems. The modularity of the lumen enables rational engineering of multi-ion traps, chemically gated constrictions, and sequence-specific reader elements, offering a path toward overcoming current resolution limits in DNA and protein sequencing. More broadly, programmable *de novo* TMBs provide a foundation for tailored selective transport, molecular sequestration, and responsive membrane devices. As generative models increasingly incorporate global barrel dynamics and membrane-aware constraints, we anticipate that β-barrel pores will evolve from structural mimics of natural proteins into fully engineerable interfaces between biological molecules and synthetic systems.

## Acknowledgements

We thank David Kim for helpful discussions and for providing the 2D strand map script, Ljubicia Mihaljević for helpful discussions, and Jason Qian for guidance on DNA experiments using the ONT rapid sequencing kit.

https://github.com/davidekim/generate2dstrandmap

This work is based upon research conducted at the Northeastern Collaborative Access Team beamlines, which are funded by the National Institute of General Medical Sciences from the National Institutes of Health (P30 GM124165). The Eiger 16M detector on the 24-ID-E beam line is funded by a NIH-ORIP HEI grant (S10OD021527). This research used beamtime awards (DOI: https://doi.org/10.46936/APS-190993/60014759) from the Advanced Photon Source, a U.S. Department of Energy (DOE) Office of Science User Facility operated for the DOE Office of Science by Argonne National Laboratory under Contract No. DE-AC02-06CH11357. This research used resources 17-ID-2 of the National Synchrotron Light Source II, a U.S. Department of Energy (DOE) Office of Science User Facility operated for the DOE Office of Science by Brookhaven National Laboratory under Contract No. DE-SC0012704. The Center for BioMolecular Structure (CBMS) is primarily supported by the National Institutes of Health, National Institute of General Medical Sciences (NIGMS) through a Center Core P30 Grant (P30GM133893), and by the DOE Office of Biological and Environmental Research (KP1605010).

## Funding

Air Force Office of Scientific Research under award number FA9550-22-1-0506 (SM)

IPD Breakthrough Fund(IBF) for “de novo pores” (AP, RS)

National Science Foundation Graduate Research Fellowship Program under DGE 2137420 (CB)

Any opinions, findings, and conclusions or recommendations expressed in this material are those of the author(s) and do not necessarily reflect the views of the National Science Foundation.

## Author Contributions

Conceptualization: DB, SM

Distillation Set Curation: RS, SM, RDK

Pipeline development: AP, RS, SM, ML

Codebase for TMB_RFD2: YL, AP, BC

Training TMB_RFD2: YL, RS, AP, SM

Codebase TMB_MPNN: FX, SM

Training TMB_MPNN: FX, SM

Design and experimental characterization: RS, AP, ML, SM

JIBE experiments: CB, SM, AM, ET

Crystal structures: AB, JM, AK

Supervision: DB, SM

Funding: DB

Writing: AP, RS, SM with support from YL, FX, RDK, DB

AP, RS, and SM conceived the study, and may change the order of their names for personal pursuits to best suit their own interests.

## Competing Interests

AP, RS, SM, ML, and DB are listed as co-inventors on a patent application filed by the University of Washington related to the proteins described in this work.

## Notes

### Summary of Updates

This version of the manuscript has been revised to include data demonstrating the insertion of designed nanopores into pure block copolymer membranes. The text has also been revised throughout to improve clarity, accuracy, and flow.

