## Supplementary Information for "Generative design of programmable asymmetric β-barrel nanopores"

**The PDF file includes:**

**Materials and Methods**
**Supplementary Text**
**Figs. S1 to S31**
**Tables S1 to S4**

### Materials and Methods

#### Curation of the distillation set

We created our distillation set by mining the "Is It A Barrel?" database for putative single-chain transmembrane β-barrel (TMBB) outer-membrane proteins and retained only sequences shorter than 600 amino acids that had complete amino acid sequences (1,938,935 sequences). To minimize redundancy while maintaining adequate sequence and structural diversity for model training, clustering was performed using sequence similarity rather than structural similarity. We clustered the filtered sequences with MMseqs2 using 70% sequence identity and an 80% alignment coverage threshold, yielding 485,379 clusters. For each cluster representative, we then ran BLASTP against the Alphafold Database (AFDB50)(*26*) to identify entries whose structures are already available. These (~14,000) were excluded so that the remaining representatives could be structure-predicted with Alphafold2 (with MSAs; ~485,000 single-chain predictions) using all five network parameter sets to account for stochastic variation among predictions. Models with mean predicted Local Distance Difference Test (pLDDT) scores below 75 were excluded to remove low-confidence structures. Native structures can have signalling peptides, flexible linkers and soluble domains on top of the barrels. In some cases, barrels can also be stacked on top of each other. Only the transmembrane barrel portion of each structure was considered for downstream filtering, as described below.

For structural filtering and annotation, we processed each predicted PDB using a Rosetta-based pipeline (https://github.com/baker-laboratory/TMB_pipeline). It first assigned a secondary structure and identified β-strand segments above a minimum length. Then, it computed backbone-backbone hydrogen bonds for those segments and built an adjacent strand-strand pairing map to ensure the structure formed a closed barrel with a single chain, thus excluding any potentially oligomeric barrels. We only chose barrels that had a minimum of 8 strands and a minimum of 4 backbone-backbone hydrogen bonds per adjacent strand pair. To isolate the barrel from extraneous domains, we then determine barrel-only residues by aligning Cα coordinates of strand residues to the barrel principal axis and trimmed residues that fall outside the median strand span along Z, effectively removing regions outside the barrel cylinder. Finally, to verify consistency with transmembrane β-barrel topology, β-strand residues were classified into pore-facing (core) and lipid-facing (surface) positions using Rosetta's LayerSelector, and designs were required to contain predominantly hydrophobic lipid-facing strand surfaces (≥60%). The final distillation set is composed of 117,447 monomeric TMBs. Although all proteins in the dataset share the defining β-barrel fold, their sequence space exhibits substantial heterogeneity in strand number, loop topology, and surface polarity distribution.

#### TMB_RFD2 fine-tuning and conditioning

The curated distillation set was used to fine-tune and condition RFdiffusion2 (RFD2). When fed into the RFD2 framework, each structure, whether derived from the distillation set or from the PDB, is converted into a series of tensorized inputs that describe both local and global geometric relationships essential for β-barrel folding.

The secondary-structure (SS) tensor is a one-dimensional tensor (N × 4) in which each residue i is represented by a one-hot vector derived from DSSP assignments: helix (H), strand (E), loop (L), or others (N).
Feature-specific binary tensors, represented as one-dimensional tensors (N × 1), are included to flag residues involved in key structural features such as β-bulges and glycine twists. Each feature type is encoded in a separate tensor, where a value of 1 indicates that a residue participates in that specific feature and 0 indicates its absence. This feature encoding was motivated by prior scaffold-guided design experiments showing that designs with explicitly placed glycine twists can yield experimentally validated crystal structures (S2).
The adjacency matrix is represented as a two-dimensional tensor (N × N), where N is the number of residues in the protein. Each element (i, j) encodes the spatial relationship between residues i and j, defined by the distance between their Cα atoms. Specifically, a token of 0 indicates that the residues are not within 8 Å of each other, 1 indicates that they are within 8 Å of each other, and 3 denotes that the residues are engaged in β-strand pairing. A token of 2 represents a masked position, meaning that the relationship between those residues (0, 1, or 3) is hidden from the model. To summarize these features, Table S1 lists both the existing RFD2 features and the new key features introduced for β-barrel design, and Supplementary Fig. S10 provides a schematic of the training and inference workflow. TMB_RFD2 was trained for 20 epochs (Fig. S5). Training initially emphasized TMB structures from the distillation set, with increasing fractions of PDB monomer and PDB interface structures introduced in subsequent rounds.
We compared inference using fully unmasked adjacency and secondary-structure (SS) tensors with a partial masking strategy applied to both tensors. Specifically, the masking strategy masked most strand-pairing residues in the adjacency matrix while retaining only a small subset of residues near the center of each strand (Fig. S11). Similarly, most residues in the secondary-structure (SS) tensor were masked except for a few central residues within each strand (Fig. S11). After benchmarking both approaches, we found that partial masking produced the highest fraction of designs that passed the geometrical filters used to classify structures as barrels (Fig. S11).

**TMB_MPNN fine-tuning and conditioning**

We extended the effective receptive field of the LigandMPNN from 32 to 96 nearest neighbors, and doubled the dimension of all the node and edge embeddings. To store this additional long-range structural information withoutover-squashing, we doubled the dimension of all the node and edge embeddings. We refer to this model as 96NN_MPNN. Starting from the 96NN_MPNN model, we fine-tuned the PDB-pretrained model on the curated TMB distillation set. This model is referred to as TMB_MPNN.
To enable sequence design conditioned on transmembrane hydrophobic thickness, an approximate hydrophobic thickness of each input was first estimated by geometric analysis of the barrel orientation and residue environment. Each barrel was aligned along the Cartesian z-axis and centered at the origin, with the N- and C-terminal ends oriented toward negative z

values. Surface-exposed β-strand residues were identified based on solvent accessibility and secondary structure assignments. A tentative hydrophobic thickness was then calculated as the average distance between aromatic-rich residue rings located on opposite ends of the barrel, or between aromatic- and arginine-rich interfacial rings where present. Residues located within this hydrophobic thickness were labeled as buried (class 2), whereas those within the interfacial aromatic or arginine-rich regions were labeled as interface (class 1). These labels were added as residue-level inputs to both encoder and decoder message-passing blocks, serving as conditioning signals in addition to the standard geometric and sequence embeddings. During inference, a desired hydrophobic thickness can be specified. The generated β-barrel is aligned along the z-axis, and surface-exposed residues within this hydrophobic span are identified starting from the cis side. By default, the central 75% of these residues are assigned the buried label (2), while the outer 12.5% at each end are assigned the interface label (1). These percentages can be modified within the codebase (https://github.com/baker-laboratory/TMB_pipeline). These labels are then provided to TMB_MPNN to guide sequence generation consistent with the target hydrophobic thickness. This checkpoint works best for designing barrels with increased hydrophobic thickness.
Additionally, a bias against serine and threonine in pore-facing residues was introduced by further tokenizing surface-facing residues as token 2 and pore-facing residues as token 0 within the buried region. This checkpoint performs best for designing barrels with conventional hydrophobic thickness compatible with lipid bilayers.

**Design of de novo TMBs**

Backbones were generated using the NC_TMB_RFD2 and TMB_RFD2 model. Required inputs include the number of β-strands and the average number of residues per strand. These inputs are used to automatically construct partially specified secondary structure and adjacency matrix tensors with a wrapper script (https://github.com/baker-laboratory/TMB_pipeline), which are provided to the diffusion model through the job submission script to guide backbone generation toward desired β-barrel architectures.
Optional inputs in TMB_RFD2 allow additional geometric control, including specifying strand positions for glycine twists and β-bulges to increase backbone curvature, defining symmetry constraints, and providing structural motifs for motif scaffolding (https://github.com/baker-laboratory/TMB_pipeline). Sampling a broader range of glycine-twist positions and combinations improved expression (Fig. S14 vs. Fig. S15) and SEC profiles (Fig. S18 vs. Fig. S19), with a higher fraction of designs exhibiting predominantly monomeric peaks compared with the more limited sampling condition (Table 1 vs. Table S3). To broadly sample twist placements, designs containing two glycine twists were generated using the pattern (i, i + k), where k ≥ 2, and designs containing four glycine twists were generated as two adjacent twist pairs using the pattern (i, i + 1, k, k + 1). To target barrel geometries such as triangular shapes, glycine twists can be placed on three approximately equidistant strands around the barrel, while square-like geometries can be obtained by specifying four evenly spaced strands. The wrapper

script (https://github.com/baker-laboratory/TMB_pipeline) then automatically places the glycine twists at the specified strand numbers using residues centered around the middle of each strand, corresponding to five residue positions: nres/2-2, nres/2 - 1,nres/2, nres/2+1, nres/2 +2, where news is the number of residues per strand. β-bulge placement can be achieved by specifying residues at positions $p_i = i \times nres - 1$ for even-numbered strands i.
When motifs are provided, the model preserves the specified motif geometry while generating the surrounding backbone. For designing TMBs with copper-binding sites, a three-residue histidine motif, including the backbone coordinates, was extracted from bovine copper-zinc superoxide dismutase (BSOD) and provided as input to TMB_RFD2.
For generation of symmetric repeat alpha-beta TMB pores from scratch, cyclic symmetry at inference was implemented in TMB_RFD2 in the same manner as RFDiffusion(*24*). Thereafter, different symmetries were sampled along with necessary inputs for regular TMB_RFD2 diffusion (C4 and C8 for 16 stranded barrel) but with no chainbreaks specified in the contigs resulting in symmetric repeat output structures. These were then designed first using TMB_MPNN, filtered with Alphafold3 and the helical domain further designed with soluble ProteinMPNN followed by another round of structure prediction by Alphafold3.

**Protein Expression and Purification**
All designs were purified from *E. coli* following a protocol adapted from Vorobieva et al. Designs were ordered as synthetic genes (eBlocks, Integrated DNA Technologies) with compatible BsaI overhangs for cloning into the target vector LM0627 via Golden Gate assembly, with a stop codon introduced prior to the His-tag in LM0627. Golden Gate subcloning reactions were performed in 96-well PCR plates in 1 µL volumes. Reaction mixtures were transformed into chemically competent *E. coli* BL21(DE3) cells, followed by recovery in 100 µL SOC medium, and then distributed into four 96-deep-well plates containing 900 µL auto-induction medium composed of 1% (w/v) tryptone, 1% (w/v) sodium chloride, 0.5% (w/v) yeast extract, 1% (w/v) kanamycin. After overnight incubation at 37°C (20-24 h), cultures were scaled up into 50 mL of autoclaved TB-II medium supplemented with 1% (w/v) kanamycin, 2 mM $MgSO_4$, and 1× 5052 for further expression.
Cells were harvested and lysed. Inclusion body pellets were washed several times with buffers containing 1% Triton X-100 and 1% Brij-35, alternating between the two detergents to remove residual lipids and other contaminants. Each washing step involved resuspension of the insoluble pellet in the appropriate buffer, brief sonication, and incubation for one hour at room temperature or overnight at 4°C.
After washing, pellets were solubilized in GuCl, and 500uL of the denatured protein solution were refolded in 30mL of a buffer containing 25 mM Tris-Cl (pH 8.0), 100 mM NaCl, and 0.1% DPC (dodecyl-phosphatidylcholine). Refolding was carried by spontaneous dilution.
The refolded samples were incubated overnight at 4°C with gentle shaking, concentrated using 10 kDa molecular-weight cutoff filters, and purified by size-exclusion chromatography (SEC) on a Superdex 200 Increase 10/300 GL (Cytiva) column. Fractions eluting at the expected volume

for the monomeric state (~99mL) of the designed pore were pooled, concentrated, and stored at 4°C for subsequent analysis.

Representative expression and purification data, including SDS-PAGE gels and SEC chromatograms, are shown for designs listed in Table 1, motif-scaffolded designs, and longer-barrel designs in Figs. S13, S14, S16–S18, and S20.

**Electrophysiological Characterization**

Membrane insertion and conductance measurements were performed using an Orbit 16 TC instrument (Nanion Technologies GmbH, Munich, Germany) on MECA chips. Lipid stock solutions were prepared in dodecane at a concentration of 1 mg/mL, using DPhPC (di-phytanoyl-phosphatidylcholine) lipids for all experiments.

Purified proteins were diluted into a buffer containing 0.05% DPC (~1× CMC), 25 mM Tris-Cl (pH 8.0), and 100 mM NaCl, to a final protein concentration of approximately 5 µg/mL. A small aliquot (~0.5 µL) of this protein solution was added to the cis chamber of the chip containing 200 µL of KCl buffer while lipid bilayers were formed using the Orbit's built-in rotating stir-bar setup. Measurements were conducted at 25°C under symmetric buffer conditions, with either 1 M KCl or 1.5 M KCl present on both sides of the membrane.

Spontaneous membrane insertion was detected as discrete current jumps characteristic of single-channel openings. Raw signals were recorded at a sampling frequency of 10 kHz and downsampled to 100 Hz using an 8-pole Bessel filter. Current traces were processed using a custom Python script (https://github.com/baker-laboratory/TMB_pipeline).

For comparison of conductance across different cationic and anionic electrolyte conditions, single-channel electrophysiology measurements were performed in symmetric 1 M electrolyte solutions containing $CaCl_2$, KCl, NaCl, LiCl, KF, KCl, and KI, and conductance values were determined from the resulting current traces.

For testing the motif-scaffolded design containing a copper-binding site within the pore (TMB_N10_HIS3_1), 2 µL aliquots of 100 mM $CuSO_4$ were added to the cis chamber three times, with incubation periods between additions to allow diffusion to the pore. Control experiments were performed under identical conditions using $CoCl_2$ and $K_2SO_4$. The same $CuSO_4$ addition protocol was also applied to the knockout design KO_TMB_N10_HIS3_1.

For testing designs in block copolymer membranes, membranes were prepared using poly(butadiene-*b*-ethylene oxide) (PBd-*b*-PEO; $Bd_9EO_5$, Mn = 0.5-*b*-0.2 × $10^3$ g/mol, PDI = 1.02).

**Circular Dichroism**

For screening designs in detergent micelles, the protein/detergent was directly analyzed by CD spectrometry in SEC buffer (25 mM Tris-Cl (pH 8.0), 150 mM NaCl, and 0.1% DPC (dodecyl-phosphatidylcholine)). CD spectra were obtained using a Jasco model J-1500 spectropolarimeter over a wavelength range of 260-190 nm at room temperature. Average CD spectra from three repeats were obtained using a Chirascan Plus (Applied Photophysics).

**ssDNA experiments**

A 22 nucleotide Thymine repeat ssDNA was purchased from IDT and used for the ssDNA translocation experiments on TMB_N14_des12. 2uL of the DNA was added on top of a single pore in a DPhPC bilayer from a 1uM stock solution in water. The bilayer was painted across a symmetric solution of 25mM Tris and 200mM NaCl in a MECA 100uM chip. The applied voltage was changed in increments of 25mV from 50 to 150 mV and the corresponding current recorded at a sampling frequency of 20kHz. The recorded signal was then filtered using a digital 8 pole low-pass bessel filter at 5kHz and simple thresholding based event detection was carried out using custom scripts in python (https://github.com/baker-laboratory/TMB_pipeline). The dwell time histogram was fitted with an exponential distribution and their corresponding means plotted at different applied voltages.

**DNA threading experiment**

The Oxford Nanopore sequencing kit (SQK-RAD114) was used following their suggested protocol for lambda phage DNA to prepare the Sequencing Buffer (SB) plus DNA mix. A custom in-house PET29b vector backbone was used instead of the lambda DNA. In a MECA 100uM 16 cavity chip, TMB_N14_des1 pore was reconstituted in DPhPC lipid bilayers across a symmetric salt solution containing 1M KCl and 25mM Tris at pH 8.0. The solution temperature was increased to 38C using the Orbit16 TC temperature controller and 40uL of the sequencing mix was added near the cavity containing the pore. The added buffer was pipetted up and down a couple of times and the applied voltage was increased to 150mV slowly in steps of 10mV increments. The current signal was recorded at 10kHz sampling frequency and subsequently filtered using a 8 pole low-pass bessel digital filter to 2kHz. The recorded current was not normalised to compare raw current perturbations to the standard ONT R10 pore signal with the same sequencing mix. For both cases, events were first estimated using the PELT algorithm with the ruptures python library. Then a histogram of all the found levels was fitted with a bayesian gaussian mixture model (GMM) with a maximum of 15 states with the sklearn library. Then using a 6 percent threshold for state weights, an optimal number of states was selected for subsequently fitting a hidden markov model (HMM) model. The hmmlearn library in python was used to fit the hmm model to the filtered signal with initialised means of PELT levels from a deterministic GMM model using the optimal number of states as determined above. The filtered signal was divided into 90:10 train:test set for the HMM training and validation. Predicted current states on the validation signal were plotted with shaded regions for each state indicating the square root of their covariance matrices.

**JIBE experiment 1 (Figure 5c)**

The monoglyceride JIBE was formulated by dissolving monolinolein (Nu-Chek Prep Inc., MN) with squalane oil (≥ 90%, Sigma Aldrich) to a concentration of 10 mg/mL. Monolinolein has a chain length of 18 carbons with two points of unsaturation at carbon 9 and 12, abbreviated as

C18:2 Δ9,12 cis. This mixture was combined with squalene oil (≥ 98%, Sigma Aldrich) and hexadecane oil (99%, Sigma Aldrich) at a volume ratio of 1:1:2, respectively, to a final volume of 0.75 mL. The aqueous phase consisted of Milli-Q water (resistivity ≥ 18.2 MΩ-cm, TOC < 50 ppb, Milli-Q IQ7000, Merck) containing 200 mM KCl (Fisher Chemical) and 30 mM Bis-Tris (ULTROL® Grade, EMD Millipore Corp), adjusted to pH7 with HCl. The aqueous solution was added dropwise to the oil phase at a 1:1 volume ratio to yield a total volume of 1.5 mL. Protein solutions (1-3 µL) were added directly to the oil-water mixture to achieve a lipid-to-protein ratio (LPR) of 1500 (TMB12_3) or 3000 (TMB_N10_des5). For the negative controls, 9.38 µL of elution buffer (TBS-DPC) was added in place of protein. The oil-water mixture was vortexed at 3000 rpm for 1 min and centrifuged at 8000 × g for 1 min. Excess oil was removed after centrifugation, yielding a jammed water-in-oil emulsion in which monoglyceride-stabilized droplets formed bilayers at droplet interfaces. A custom three-part device was fabricated using a Bambu Lab P1P 3D printer with 0.2 mm PLA filament to support the JIBE during voltage-clamp experiments. The assembled device consisted of two outer plates and a central spacer (2 mm thickness) that defined the JIBE chamber. Each outer plate contained a circular aperture with an outer diameter of 10 mm and an inner diameter of 7.5 mm. The apertures were filled with 2% (w/v) agarose hydrogel (Molecular Biology Grade, Fisher Chemical) to a thickness of 2.5 mm. The JIBE was loaded into the central chamber between the hydrogel layers, resulting in a membrane area of approximately 0.78 $cm^2$. The inner diameter of the hydrogel layers is reduced compared to the outer diameter to prevent the agarose layers from being displaced out of the plates. Fluid reservoirs were secured on both sides of the hydrogel-JIBE-hydrogel assembly. Each reservoir was filled with 2.5 mL of the same aqueous solution used for JIBE preparation (200 mM KCl, 30 mM Bis-Tris, pH7).cElectrical measurements were performed using a patch clamp amplifier (HEKA EPC 10 USB) in a two-electrode configuration. Ag/AgCl electrodes (0.25 mm diameter) were placed in each reservoir on the opposite side of the JIBE membrane, with one connected to the amplifier headstage and the other grounded. To promote protein insertion into the droplet interface bilayers, a constant DC conditioning voltage was applied prior to measurement. Specifically, +1 V was applied for 15 min, followed immediately by -1V for 15 min. This conditioning step resulted in a gradual increase in current over the 30 min period, consistent with increased protein incorporation into the membrane. Omitting this step resulted in significantly lower currents during subsequent measurements. Following conditioning, current-voltage (I-V) measurements were performed by applying a DC voltage sweep from -200 mV to +200 mV in 40 mV increments. Each voltage step was held for 30 s, and current was recorded at a sampling rate of 50 kHz (20 µs) with a 1 kHz Bessel filter applied during acquisition. Membrane conductance was determined from the slope of the linear fit to the I-V relationship.

**JIBE experiment 2 (Figure 5d)**

The oil phase was prepared by adding 125 µL squalene, 125 µL 30 mg/mL C18 glycerol monolinoleate lipids in squalene, and 250 µL hexadecane to a 1.5 mL centrifuge tube. The aqueous phase was prepared as a 1 mM Tris-Cl (pH 8.0) and 10 mM KCl solution. Next, 240 µL of the aqueous phase and 10 µL of 1 mg/mL protein of interest were simultaneously added to the oil phase. The emulsion was immediately vortexed at 3200 rpm for 5 min (Scientific Industries Vortex-Genie 2), then centrifuged at 5000 rcf for 5 min (Eppendorf™ Centrifuge 5430 R) to form the JIBE.

To assemble the device, a 0.5% agarose solution was prepared with 10 mM KCl and 1 mM Tris-Cl (pH 8.0). Immediately before assembly, the temperature of the agarose solutions was 45-55°C. Using a syringe with a removable needle head, we filled the plastic chamber wells (Figure S31a) with the agarose solution, then inserted the syringe into the channel and flipped the chamber, releasing the solution into the O-ring and tilting to spread agarose thinly and evenly (approximately 1 mm thickness). After letting agarose dry for approximately 30 seconds, the syringe needle was partially untwisted, the chamber flipped over, and remaining agarose pressed out to fill the top of the needle head. Using Ag/AgCl electrodes (0.015" diameter) and an ammeter with pico to nanoampere resolution, the chambers were checked for electrical connection throughout by inserting one electrode into the agarose filled needle and gently touching the other electrode to the agarose surface within the O-ring, ensuring the current exceeded the upper current threshold (e.g. 200 nA). We pipetted the clear oil phase off the top of the JIBEs (Fig. S31b), cut off the top half of the centrifuge tube, and used flat tip tweezers to wipe JIBEs onto the surfaces of the agarose pads and O-rings on both chambers (Fig.S31c). The device was assembled by placing the plastic chambers into the larger metal chambers with the JIBE layers facing inside, then screwing the metal chamber pieces together. The electrodes were inserted into the needle heads of the plastic chambers (Fig. S31d) and a voltage applied to measure conductance. To facilitate protein insertion, a constant voltage of 450 mV was applied.

**Crystallography**

All crystallization experiments were conducted using the sitting drop vapor diffusion method. Crystallization trials were set up in 200 nL drops using the 96-well plate format at 20 ˚C. Crystallization plates were set up using a Mosquito LCP from SPT Labtech, then imaged using UVEX microscopes and UVEX PS-256 from JAN Scientific. Diffraction quality crystals formed in 0.05 M Glycine, pH 9, and 55% (v/v) PEG 400 for TMB_N10_des2; in 0.2 M Ammonium chloride, 40% (v/v) MPD for TMB_N10_des3; and 0.08 M Sodium citrate tribasic dihydrate pH 5.6, 28% v/v tert-Butanol, 20% v/v Glycerol for GlyTwistScaffold_des1.

Diffraction data was collected at the Advanced Photon Source beamline on 24-ID-E and National Synchrotron Light Source II on FMX. X-ray intensities and data reduction were evaluated and integrated using XDS(*56*) and merged/scaled using Pointless/Aimless in the CCP4 program suite(*57*). Structure determination and refinement starting phases were obtained by molecular replacement using Phaser for TMB_N10_des2(*58*) and DIMPLE for TMB_N10_des3(*59*) using the designed model for the structures. Following molecular replacement, TMB_N10_des2 and GlyTwistScaffold_des1 were improved using phenix.autobuild(*60*); efforts were made to reduce model bias by setting rebuild-in-place to false, and using simulated annealing and prime-and-switch phasing. TMB_N10_des2 was refined in Phenix(*60*). Model building was performed using COOT(*61*). The final model was evaluated using MolProbity(*62*). For TMB_N10_des3, iterative model building and corrections were performed manually using COOT(*61*) following molecular replacement and subsequent structure refinement was performed with CCP4 Refmac5(*63*). Initial refinement was conducted using BUSTER(*64*) to rapidly fix Ramachandran, rotamer, and density fit outliers, refining to convergence. PDB-REDO(*65*) was used to assess the model quality in between refinements and to fix any rotamer and density fit outliers. Final refinement was carried out in Phenix(*60*), with manual model building performed in COOT(*61*). The model quality was evaluated using MolProbity(*62*). Data collection and refinement statistics are recorded in Table S4. Data deposition, atomic coordinates, and structure factors reported in this paper have been deposited in the Protein Data Bank (PDB), http://www.rcsb.org/ with accession code **pdb_000036HN, pdb_000036HS** and **pdb_000036HO**

| Features | Dimensions | Description |
|---|---|---|
| **1D Features** | | |
| is_$\beta$-bulge | 2 | Binary indicator assigned to a single residue, encoding the presence or absence of a $\beta$-bulge at that position. |
| is_glycine_kink | 2 | Binary indicator encoding a glycine-mediated twist motif defined over a five-residue local backbone window. |
| secondary_structures | 5 | One-hot encoding with five states: helix (0), strand (1), loop (2), mask (3), and small molecule (4). |
| motif_sequence | 79 | Amino acid identity representation of the scaffolded motif sequence. |
| **2D Features** | | |
| adjacency_matrix | 4 | Pairwise residue–residue encoding with four states: far (0), close (1), mask (2), and $\beta$-strand pair (3). |
| **3D Features** | | |
| motif_structure | 3 | Three-dimensional Cartesian coordinates of motif residues, extracted exclusively from pore-facing regions during training. |

**Table S1. Input feature representations used for training TMB_RFD2.**

**Table S2. Sequences of all experimentally tested designs that exhibited conductance.**

| Designs from TMB_RFD2 (less glycine twist positions sampled) | | | | |
|---|---|---|---|---|
| **Barrel type** | **Good SEC** | **Good SEC (%)** | **Functional** | **Functional (%)** |
| N10 | 5/12 | 41.7% | 2/12 | 16.7% |
| N12 | 2/16 | 12.5% | 1/16 | 6.3% |
| N14 | 2/10 | 20.0% | 1/10 | 10.0% |

**Table S3. SEC and functional success rates for TMB_RFD2 designs with reduced glycine-twist sampling.** Percent good SEC and percent functional designs are shown for a design campaign in which glycine twists were specified on two strands positioned approximately half a barrel apart. For each barrel type, good SEC counts are reported relative to the total number of designs tested, and functional success rates are calculated relative to designs with good SEC profiles.

| | TMB_N10_des2 (PDB ID: pdb_000036HS) | TMB_N10_des3 (PDB ID: pdb_000036HO) | GlyTwistScaffold_des1 (PDB ID: pdb_000036HN) |
|---|---|---|---|
| **Data collection** | | | |
| Space group | *C 2 2 21* | *P 41 2 2* | *P 64* |
| Cell dimensions | | | |
| *a, b, c (Å)* | 87.64, 87.64, 142.50 | 70.19, 70.19, 220.67 | 86.31, 86.31, 82.21 |
| α, β, γ (°) | 90, 90, 90 | 90, 90, 90 | 90, 90, 120 |
| Resolution (Å) | 71.25 - 4.00 (4.47 - 4.00) | 34.66 - 4.00 (4.47 - 4.00) | 38.21 - 3.80 (4.79 - 3.80) |
| *Rmerge* | 2.03 (3.48) | 0.38 (0.46) | 0.432 (1.027) |
| *I / σI* | 5.7 (2.7) | 10.1 (8.2) | 6.02 (2.95) |
| Completeness (%) | 99.8 (99.8) | 99.9 (100) | 99.97 (100.00) |
| Redundancy | 11.9 (11.5) | 24.4 (23.8) | 15.7 (15.4) |
| CC1/2 | 0.821 (0.556) | 0.972 (0.953) | 0.989 (0.882) |
| **Refinement** | | | |
| Resolution (Å) | 46.76 - 4.0 (4.40 - 4.00) | 9.99 - 4.0 (4.13 - 4.00) | 38.21 - 3.80 (4.79 - 3.80) |
| No. reflections | 4843 (1202) | 4758 (477) | 3492 (1738) |
| *Rwork / Rfree* | 0.2887 (0.2510) /0.3219 (0.2508) | 0.3573 (0.3529) /0.3241 (0.3633) | 0.2450 (0.2713) /0.2894 (0.2890) |
| No. atoms | | | |
| Protein | 2424 | 3464 | 1387 |
| Ligand/ion | n/a | n/a | n/a |
| Water | n/a | n/a | n/a |
| *B-factors* | | | |
| Protein | 57 | 56 | 107 |
| Ligand/ion | n/a | n/a | n/a |
| Water | n/a | n/a | n/a |
| R.m.s. deviations | | | |
| Bond lengths (Å) | 0.002 | 0.003 | 0.004 |
| Bond angles (°) | 0.723 | 0.72 | 0.81 |

**Table S4. Crystallographic data collection and refinement statistics.** Single Crystal used for each data collection. Values in parentheses are for the highest-resolution shell.

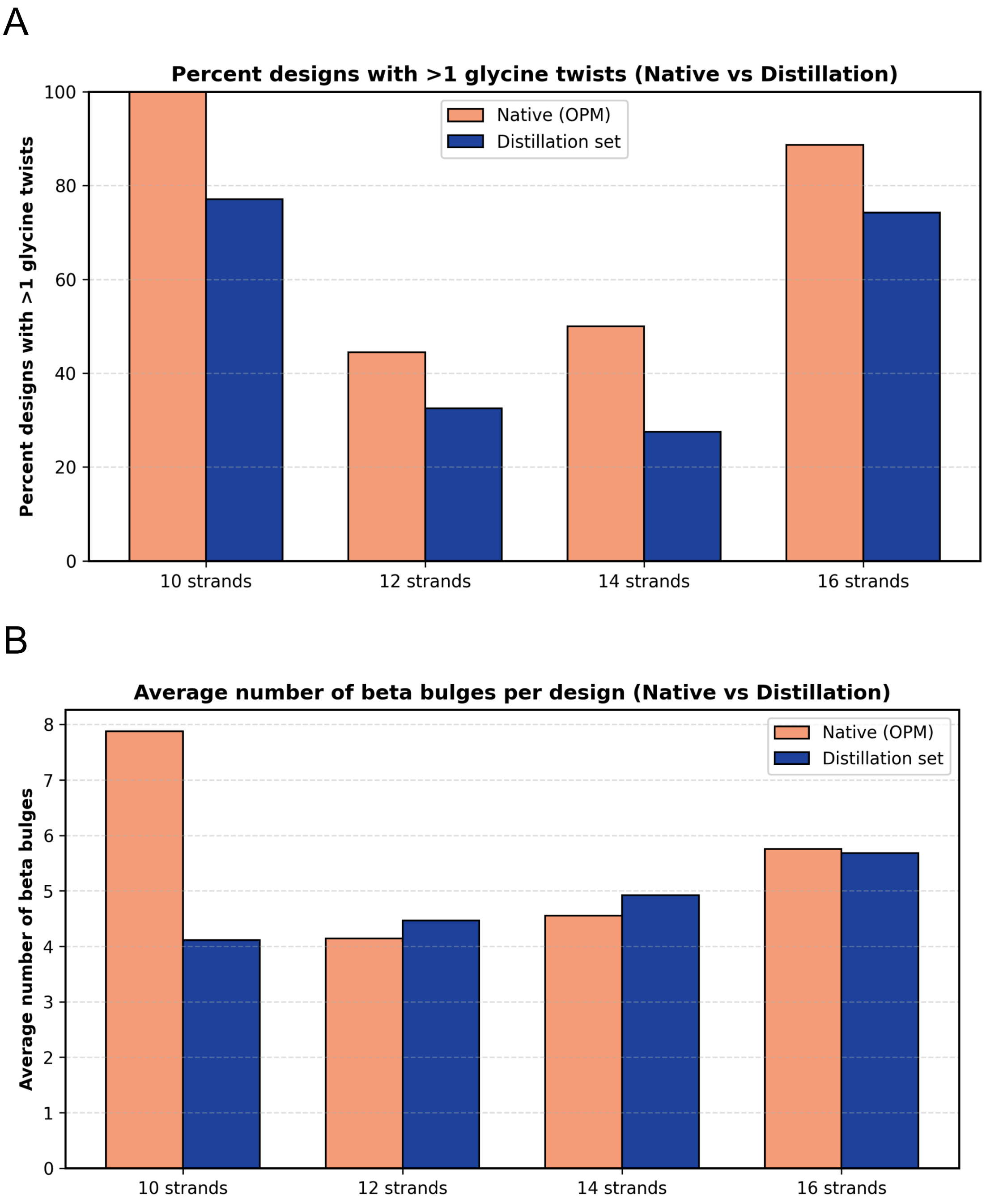


**Figure S1. Comparison of glycine twists and β-bulges in our curated distillation set and native structures from the OPM database. (A)** Comparison of the percentage of proteins with more than one glycine twist in 10-, 12-, 14-, and 16-stranded TMBs. **(B)** Comparison of the average number of β-bulges in 10-, 12-, 14-, and 16-stranded TMBs.

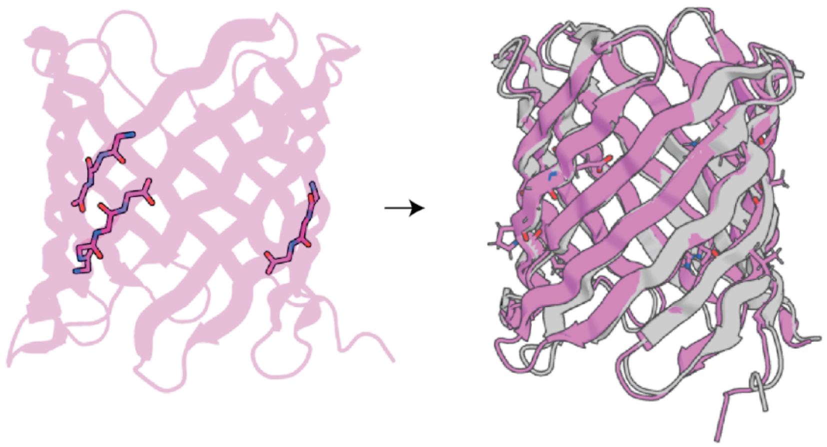

**Figure S2. Crystal structure validation of GlyTwistScaffold_des1.** The input motif is shown on the left in pink. The designed model is shown on the right in pink, overlaid with the crystal structure in gray.

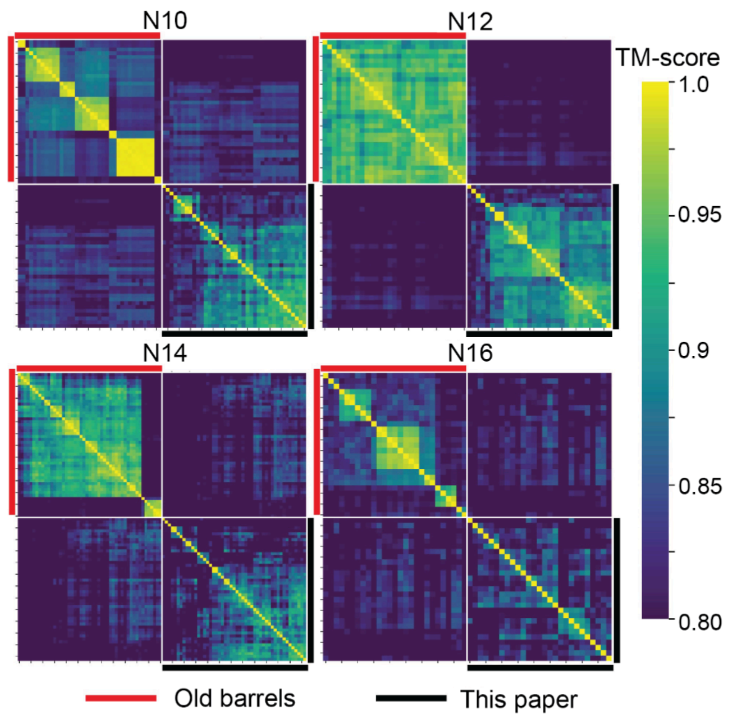


**Figure S3. TMB_RFD2 generates more structurally diverse β-barrel backbones than a parametrically guided approach.** All-by-all pairwise TM-score heatmap comparing 50 β-barrels generated using a parametrically guided approach ("Old barrels") and 50 TMB_RFD2 designs from this study ("This paper") across four strand numbers. High-TM-score clusters appear as large yellow diagonal blocks and indicate lower structural diversity within the design set.

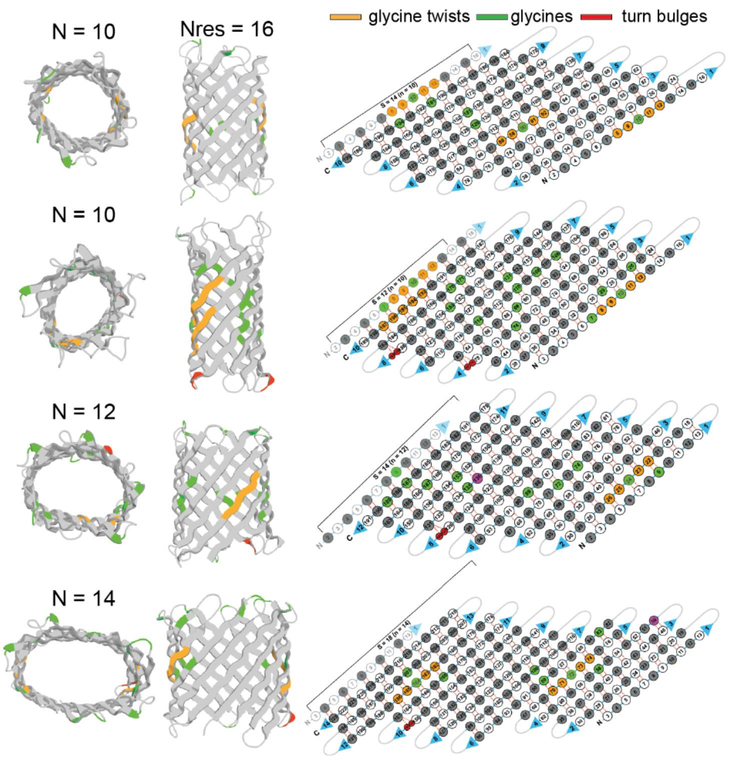


**Figure S4. Minimal input conditioning enables β-barrel generation across diverse strand shear values.** Example outputs of four β-barrels spanning a wide range of strand shear values using only two inputs: strand number (n) and residues per strand (nres). Necklace plots show diverse β-sheet registries and barrel geometries.

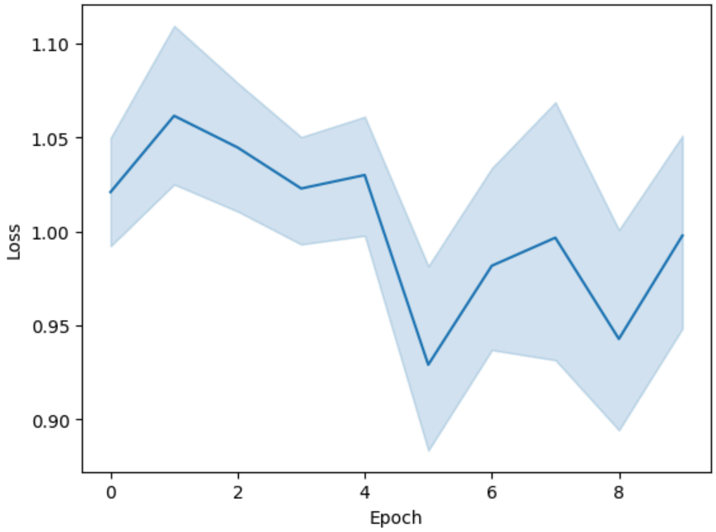


**Figure S5. Average training loss per epoch during training of TMB_RFD2.** Each epoch comprised 24,000 structures with a batch size of 8.

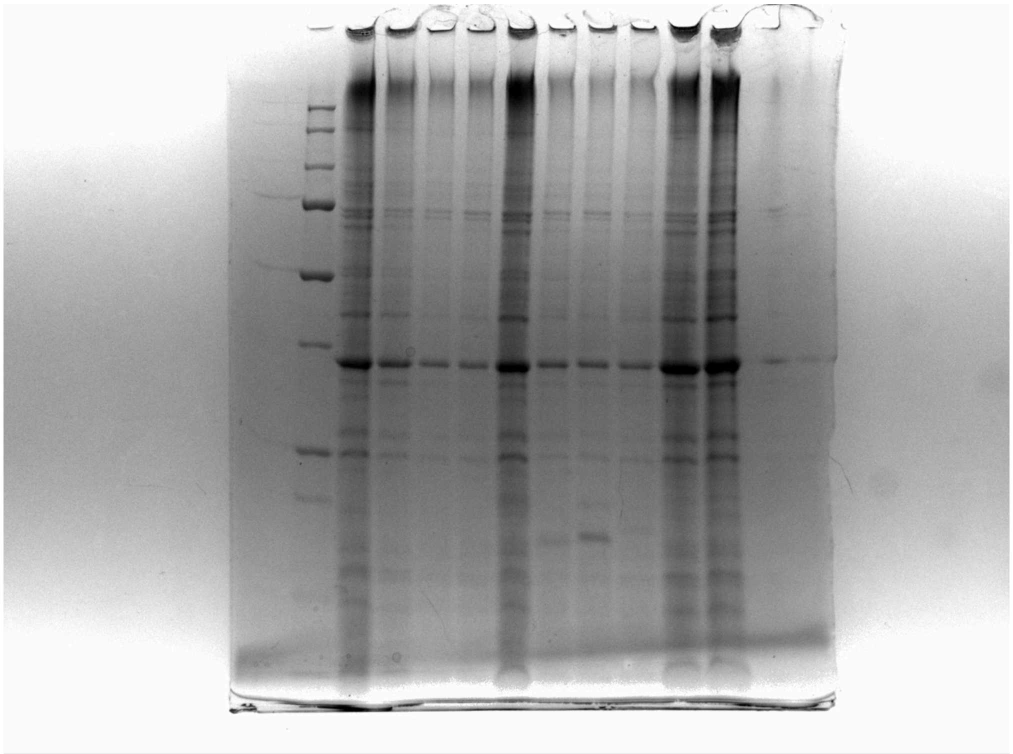

**Figure S6. ProteinMPNN-designed sequences show no expression in E. coli.** Experimental characterization of sequences designed using ProteinMPNN revealed no detectable expression in E. coli.

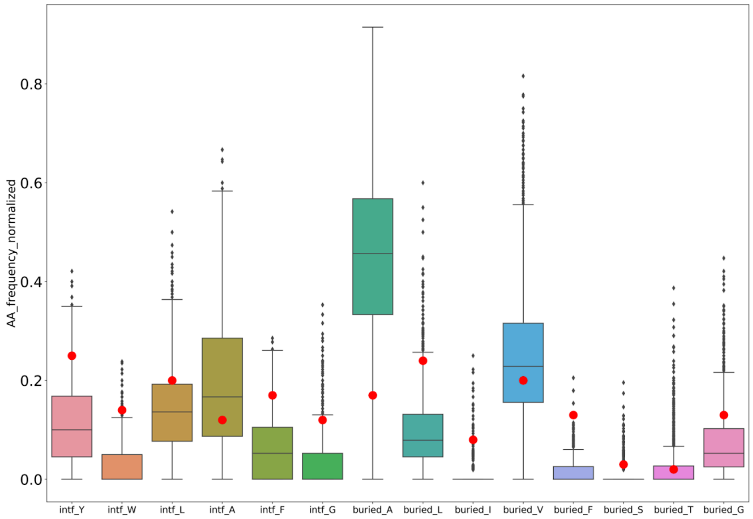


**Figure S7. Standard ProteinMPNN over-enriches hydrophobic residues in the buried regions of TMB designs.** Amino-acid distributions are shown for buried positions in TMB designs generated using standard ProteinMPNN. Red dots indicate the native TMB distribution.

A

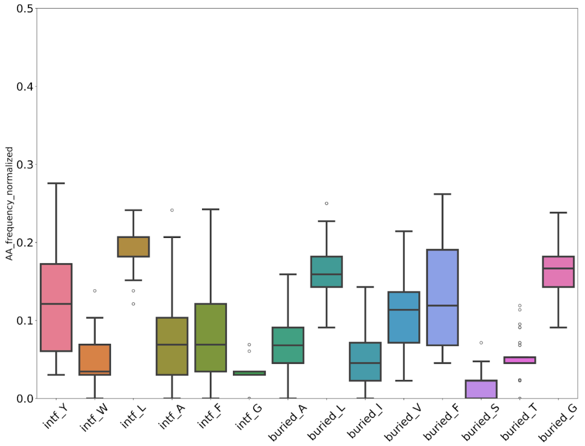


B

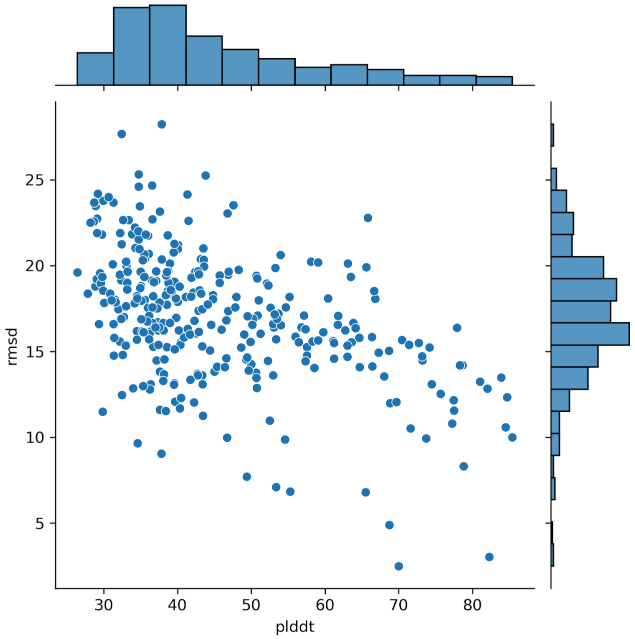


**Figure S8. Fine-tuning ProteinMPNN improves distribution of amino acids in buried region but does not reliably produce foldable TMB sequences. (A)** More uniform distribution of hydrophobics in the buried region of TMB structures with finetuned ProteinMPNN on the TMB distillation set. **(B)** Very few predicted barrels using Alphafold for the above mentioned sequences.

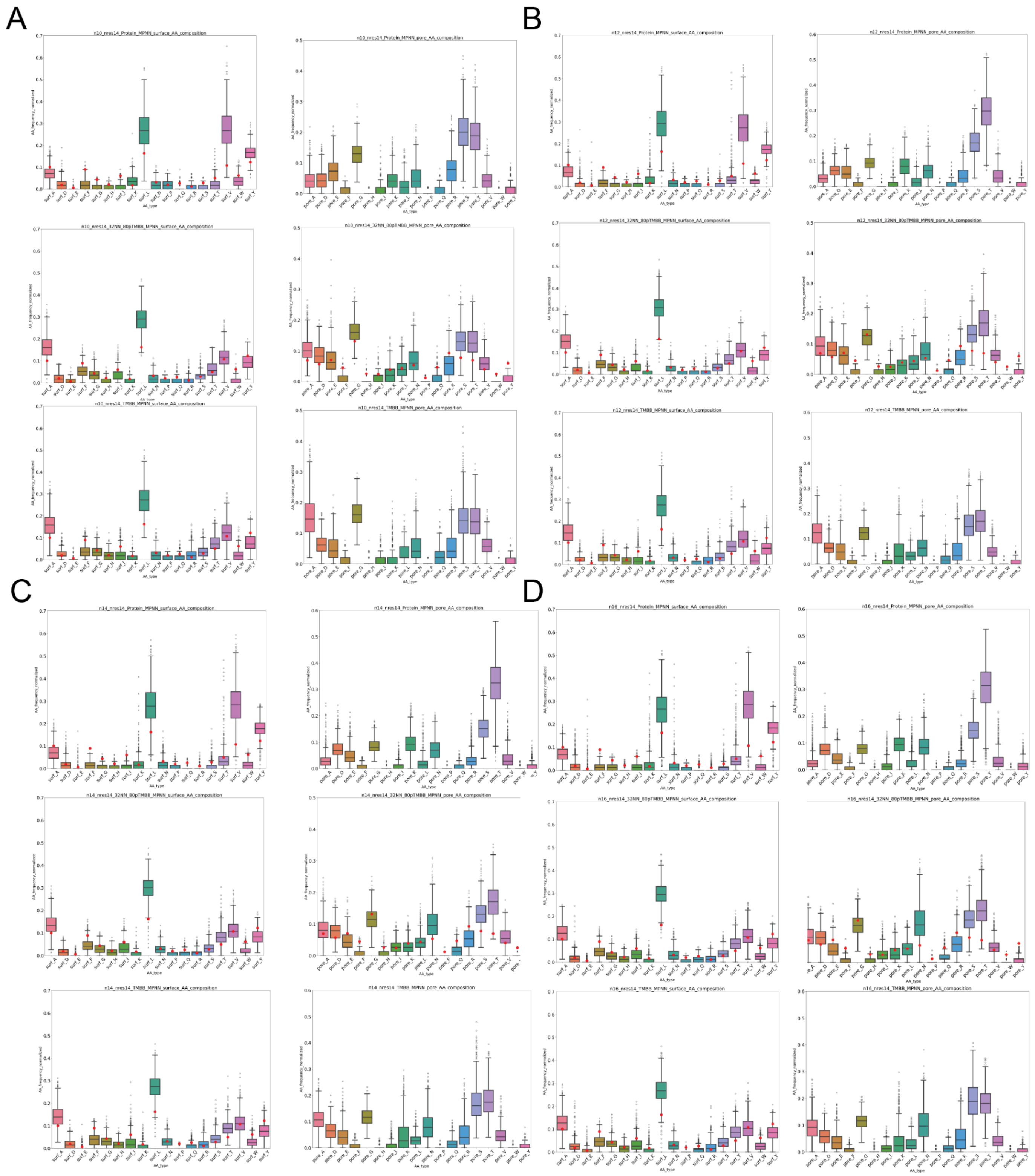


**Figure S9. Comparison of pore and surface amino acids across MPNN models. (A)** n10 **(B)** n12 **(C)** n14 **(D)** n16. The red dots show the average pore and surface amino acid composition for native barrels.

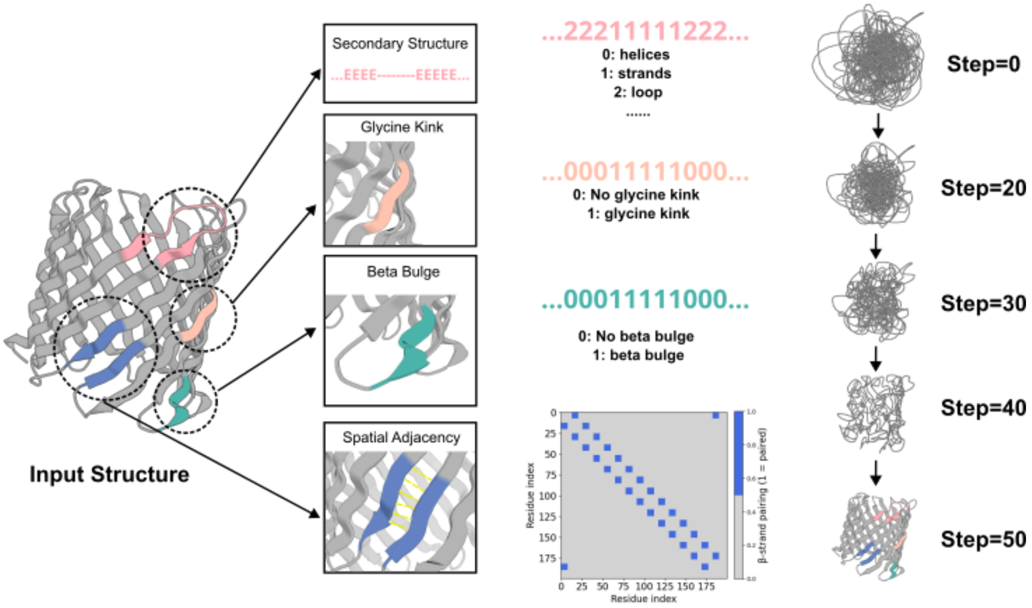


**Figure S10. Schematic representation of TMB_RFD2 training features and inference.** The model is conditioned on structural constraints including residue-wise secondary-structure assignments, glycine twists, β-bulges, and a spatial adjacency matrix defining the global β-strand pairing topology. During inference, the model starts from a disordered state and progressively refines the backbone through 50 denoising steps, converging on a folded β-barrel structure that satisfies the specified architectural constraints.

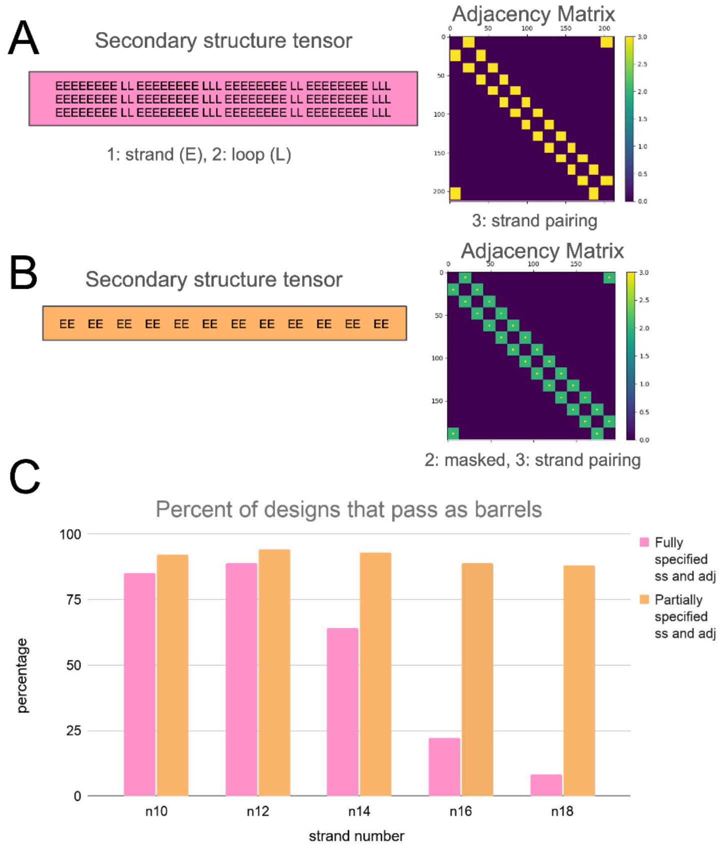


**Figure S11. Partial specification of secondary-structure and adjacency inputs improves β-barrel recovery. (A)** Example fully specified secondary-structure tensor and adjacency matrix for a 10-stranded barrel. **(B)** Example partially specified secondary-structure tensor and adjacency matrix for a 10-stranded barrel. **(C)** Comparison of β-barrel filter pass rates for designs generated using fully specified versus partially specified secondary-structure tensors and adjacency matrices. Partial specification increases the percentage of designs passing β-barrel filters across all strand numbers.

96NN_MPNN and TMB_RFD2

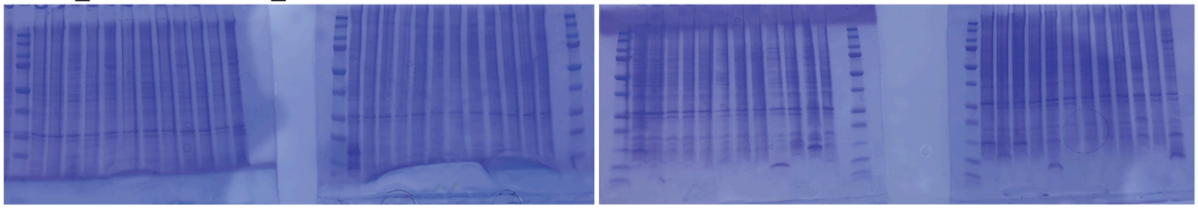

**Figure S12. SDS-PAGE analysis of TMB_RFD2 designs with 96NN_MPNN-designed sequences showing no visible expression in E. coli.**

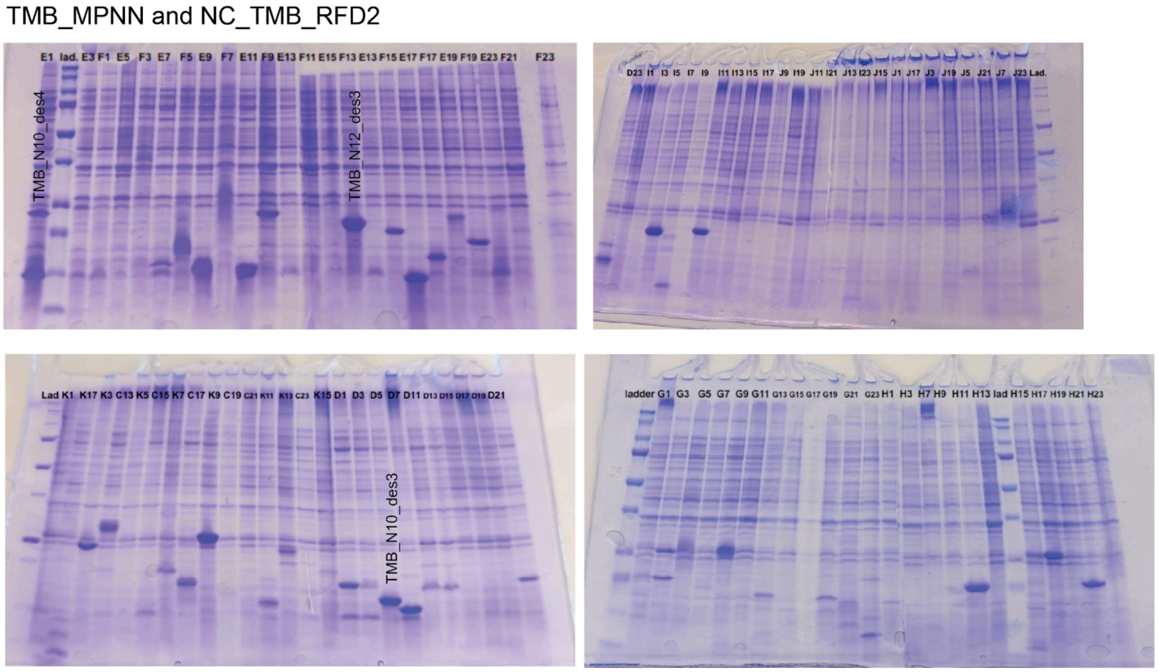


**Figure S13. SDS-PAGE analysis of NC_TMB_RFD2 designs with TMB_MPNN-designed sequences.** Functional designs are labeled according to Supplementary Table 2.

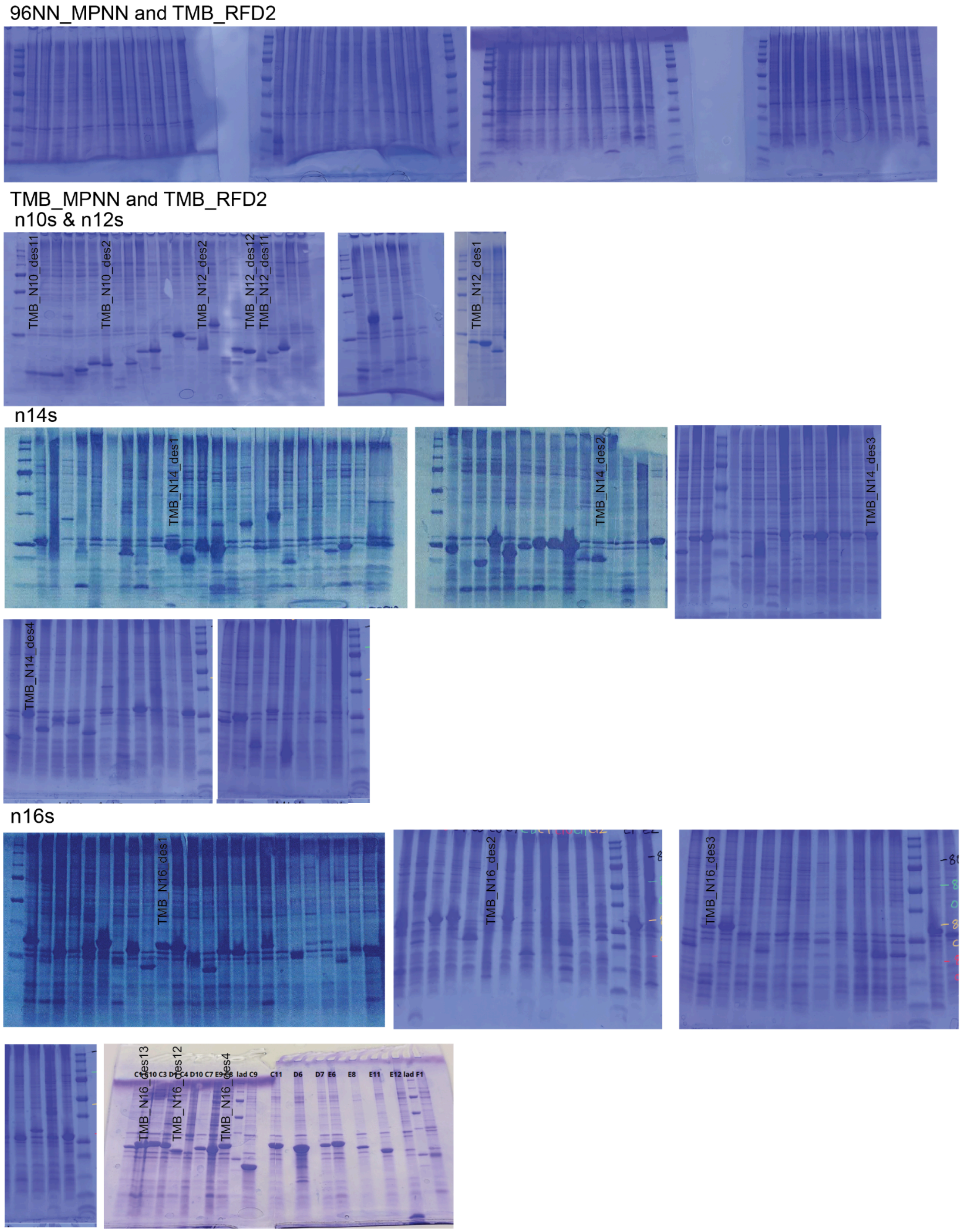


**Figure S14. SDS-PAGE analysis of TMB_RFD2 designs with TMB_MPNN-designed sequences.** Functional designs are labeled according to Supplementary Table 2.

TMB_MPNN and TMB_RFD2 (less glycine twist positions sampled)

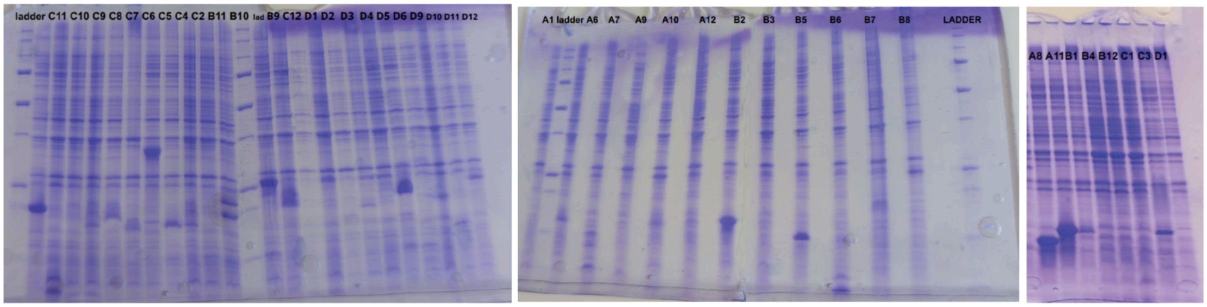


**Figure S15. SDS-PAGE analysis of TMB_RFD2 designs, generated using a reduced set of sampled glycine-twist positions, with TMB_MPNN-designed sequences.**

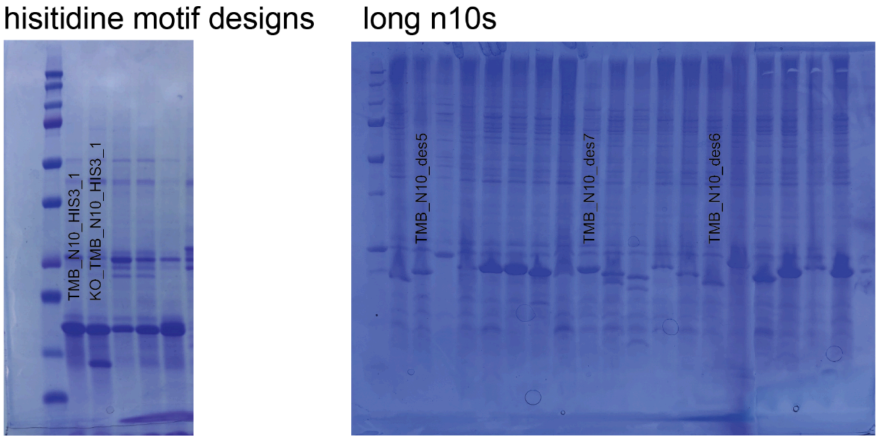


**Figure S16. SDS-PAGE analysis of motif-scaffolded and longer-barrel designs.** SDS-PAGE gels are shown for a histidine-motif-scaffolded design (TMB_N10_HIS3_1) and its knockout (KO_TMB_N10_HIS3_1), as well as for longer TMB_RFD2 backbones designed with TMB_MPNN using conditioned hydrophobic thicknesses.

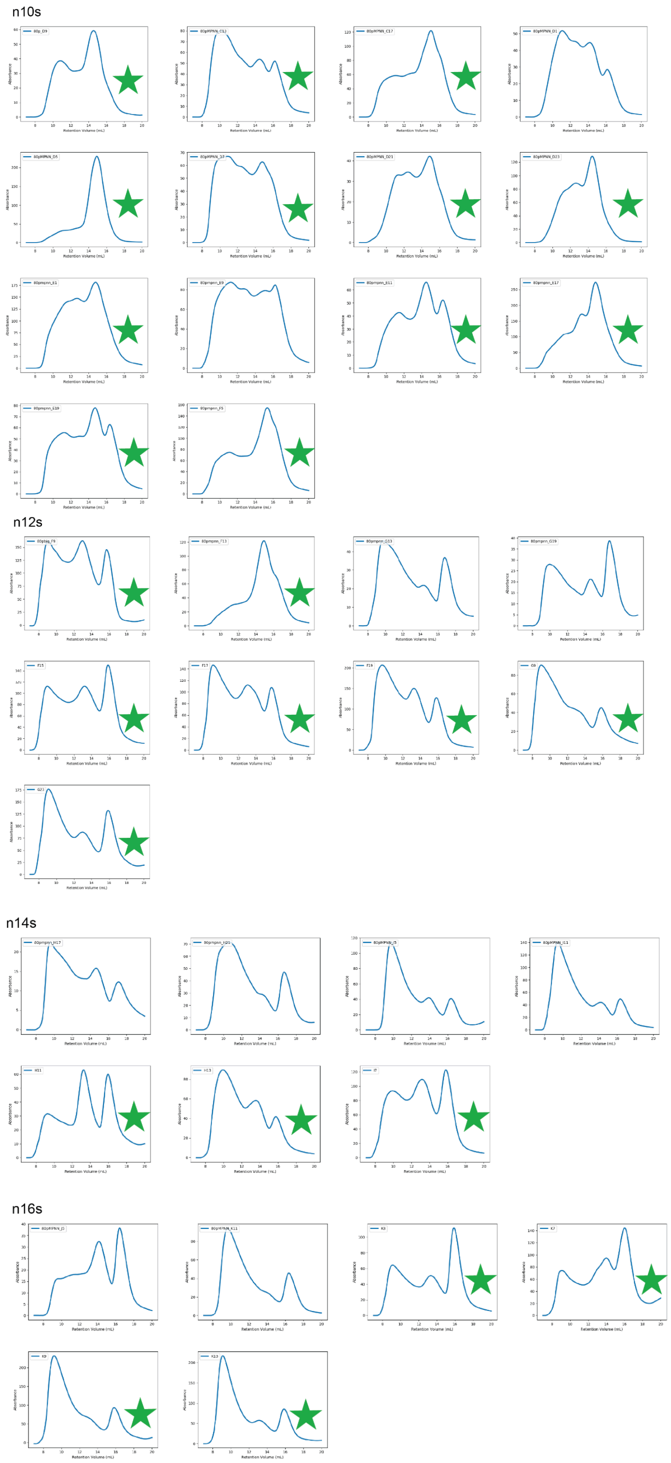


**Figure S17. SEC analysis of NC_TMB_RFD2 designs with TMB_MPNN-designed sequences.** Constructs with favorable SEC traces, marked by green stars, were advanced for characterization using the planar membrane-insertion assay and used to generate Table 1.

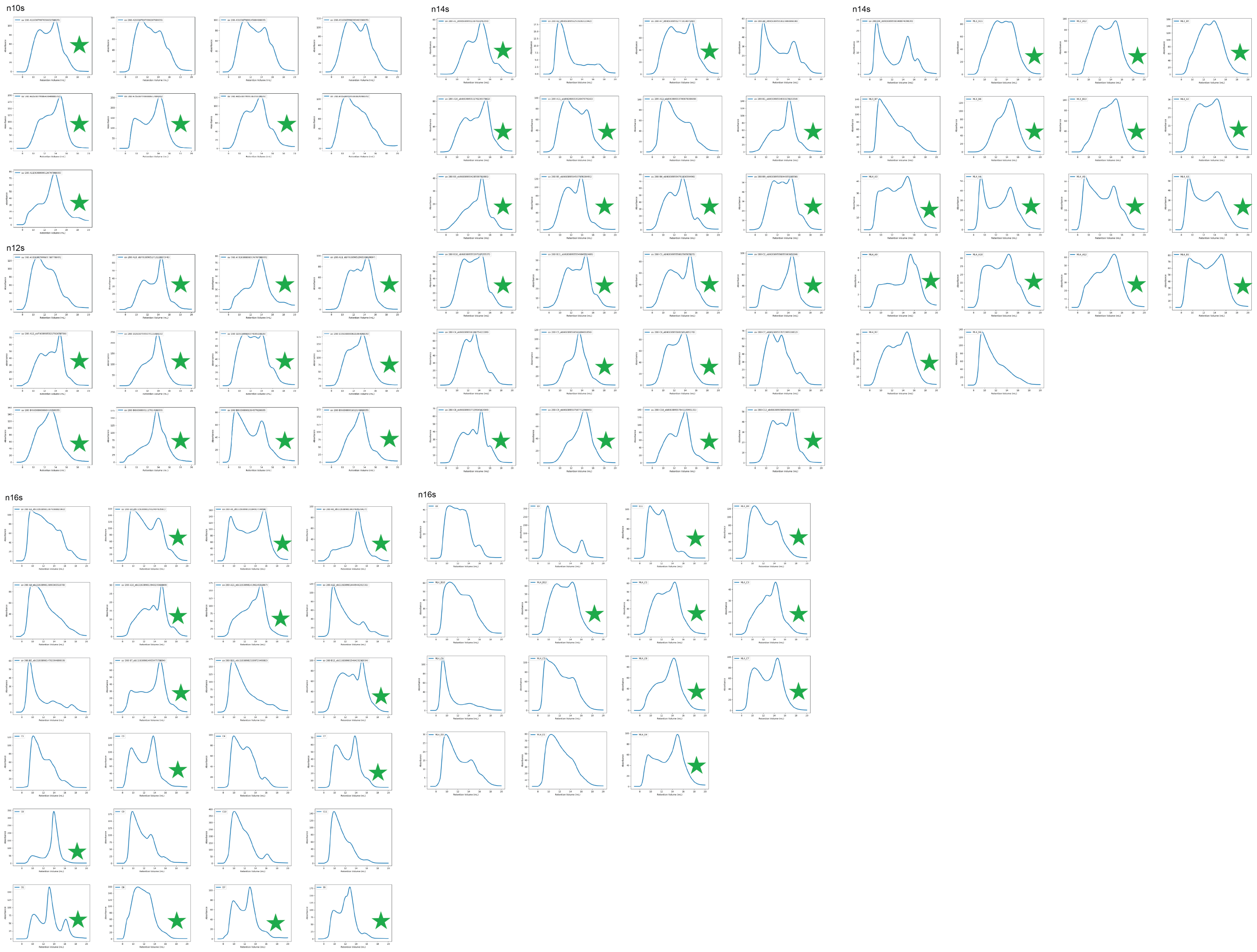


**Figure S18. SEC analysis of TMB_RFD2 designs with TMB_MPNN-designed sequences.** Constructs with favorable SEC traces, marked by green stars, were advanced for characterization using the planar membrane-insertion assay and used to generate Table 1.

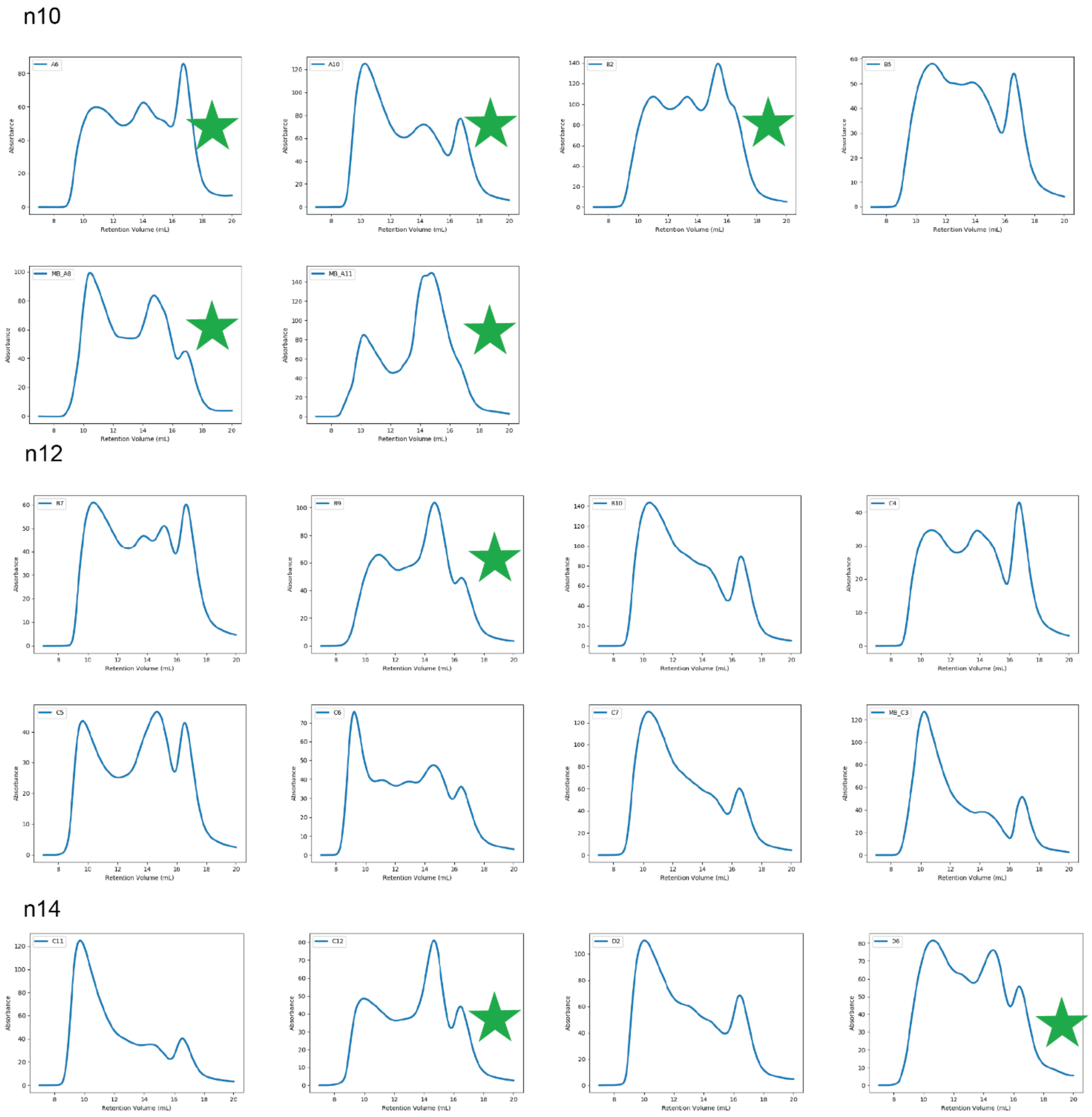


**Figure S19. SEC analysis of TMB_RFD2 designs generated with limited glycine-twist sampling and TMB_MPNN-designed sequences.** Constructs with favorable SEC traces, marked by green stars, were advanced for characterization using the planar membrane-insertion assay and used to compile Supplementary Table 3.

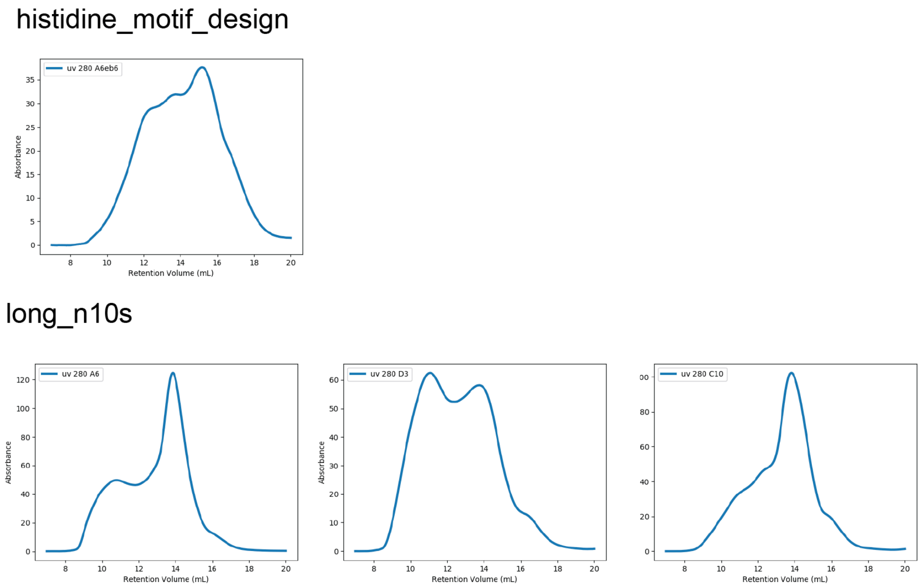


**Figure S20. SEC analysis of the histidine-motif design and longer-barrel designs.** SEC traces are shown for the histidine-motif design presented in Fig. 3 and longer-barrel designs presented in Fig. 5.

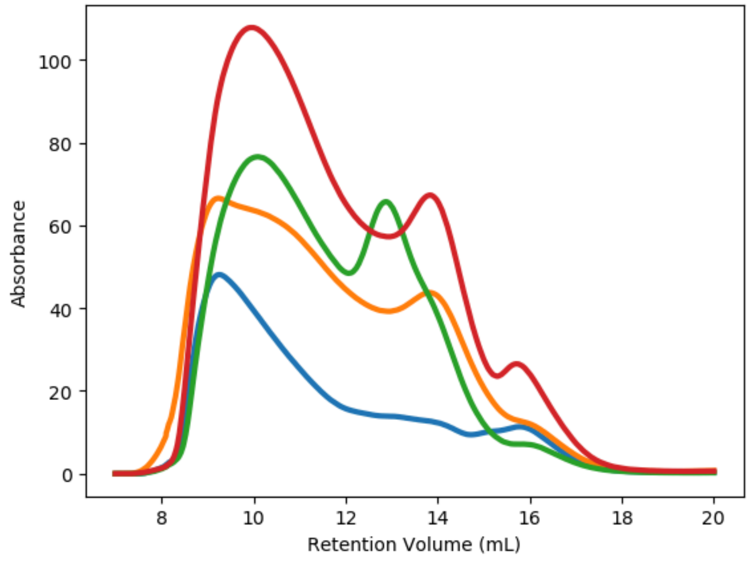


**Figure S21. SEC analysis of 18-stranded TMB designs.** SEC traces are shown for designed 18-stranded TMB backbones generated using TMB_RFD2 with TMB_MPNN-designed sequences.

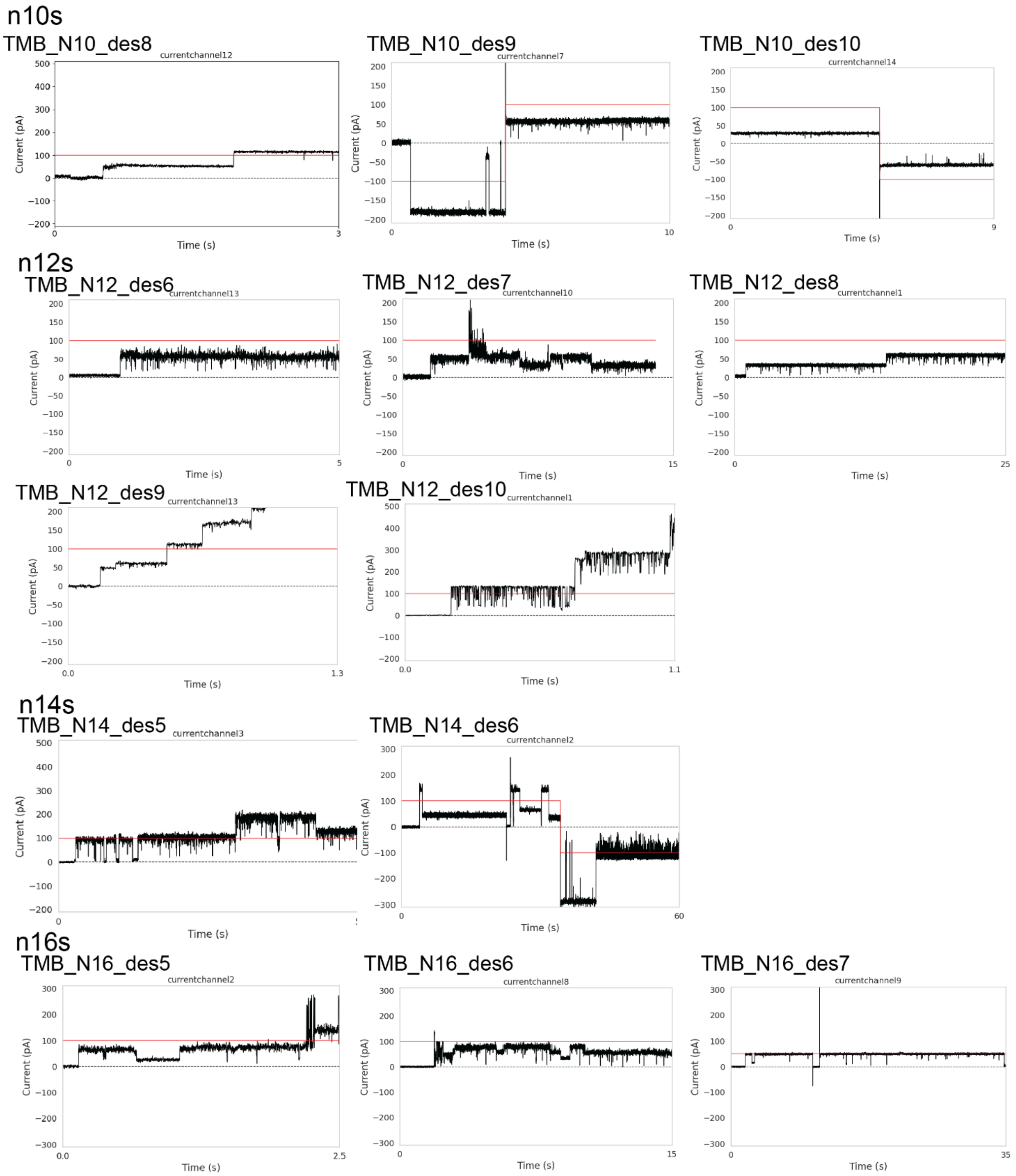


**Figure S22. Additional functional NC_TMB_RFD2 designs with TMB_MPNN-designed sequences.** Conductances are shown for functional designs not included in Fig. 3, measured in 1 M KCl in planar lipid bilayers.

A

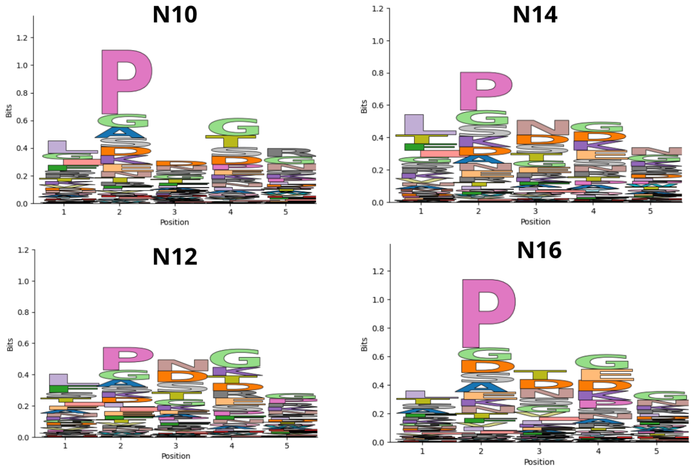
N10
N14
N12
N16
Bits
Position

B

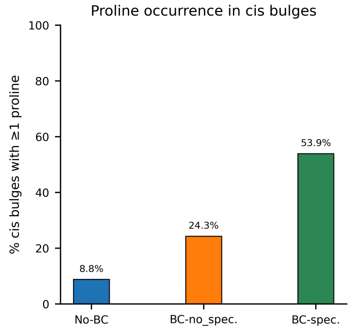
Proline occurrence in cis bulges
% cis bulges with ≥1 proline
8.8%
24.3%
53.9%
No-BC
BC-no_spec.
BC-spec.

C

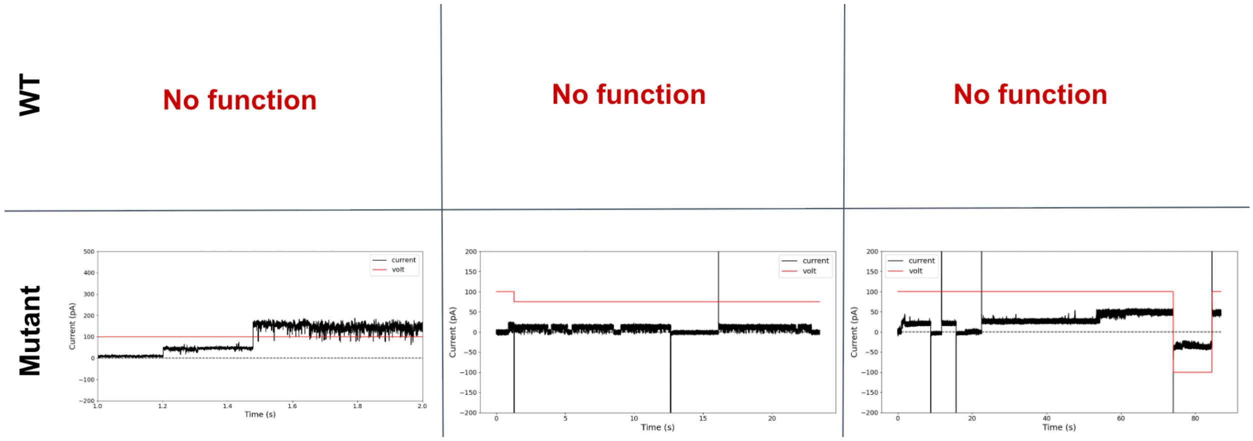
WT
Mutant
No function
No function
No function
current
volt
Current (pA)
Time (s)

**Figure S23. Prolines at cis β-bulges improve pore formation. (A)** Sequence logo plots show that proline is enriched at the second position of cis β-bulges across 10-, 12-, 14-, and 16-stranded TMBs in the distillation set. **(B)** The plot shows the percentage of cis bulges with more than one proline for various RFD2 backbone generation models showing that bulge conditioning (BC) during backbone generation increases the likelihood of MPNN placing a proline on the bulge. The No-BC model was not conditioned on β-bulges. BC-no_spec. Is the model that was trained using bulge conditioning but bulges were not specified at inference. BC-spec. explicitly specified β-bulges during inference, which further enriches proline incorporation at these sites. **(C)** Planar membrane-insertion assays show that point mutations introducing prolines at cis β-bulge positions convert previously nonfunctional designs into functional pores capable of conducting ionic current in 1 M KCl.

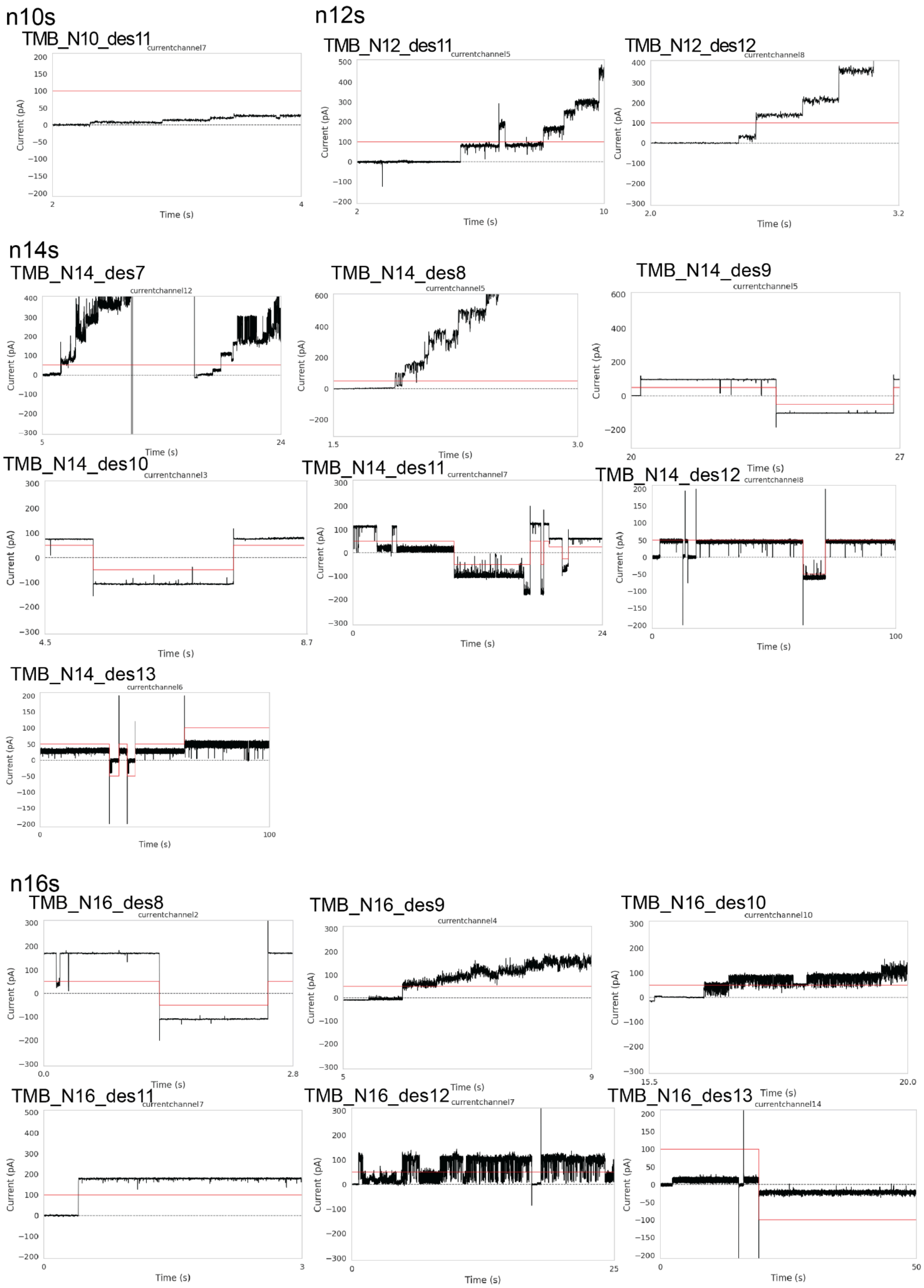


**Figure S24. Additional functional TMB_RFD2 designs with TMB_MPNN-designed sequences.** Conductances are shown for functional designs not included in Fig. 3, measured in 1.5 M KCl for 10- and 12-stranded designs and 1 M KCl for 14- and 16-stranded designs in planar lipid bilayers.

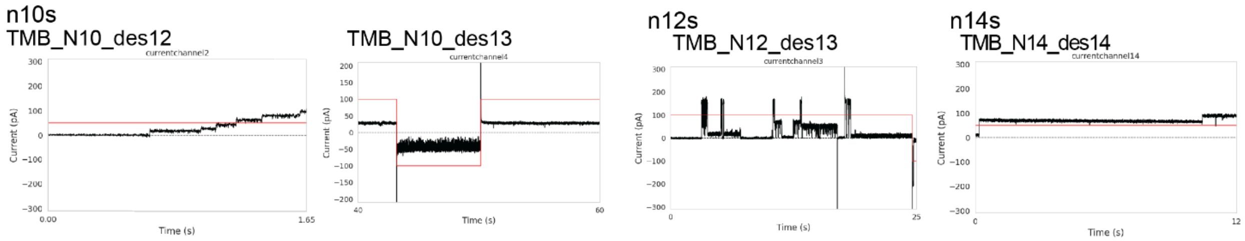


**Figure S25. Functional TMB_RFD2 designs generated with reduced glycine-twist sampling and TMB_MPNN-designed sequences.** Conductances are shown for functional designs measured in 1 M KCl in planar lipid bilayers.

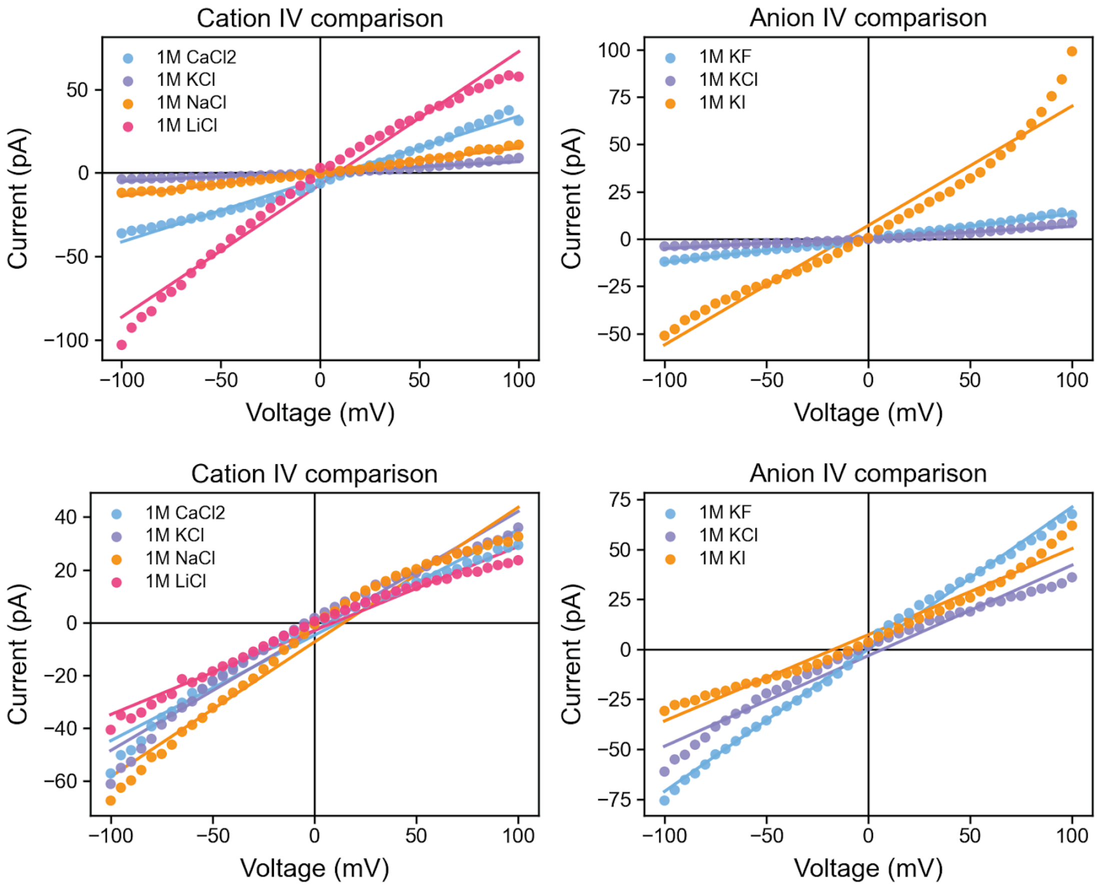


**Figure S26. Single-channel conductance of 10- and 12-stranded TMB designs under different ionic conditions.** Conductance values derived from single-channel recordings are shown for the 10- and 12-stranded designs presented in Fig. 3b, measured with different cations and anions using the planar lipid bilayer assay.

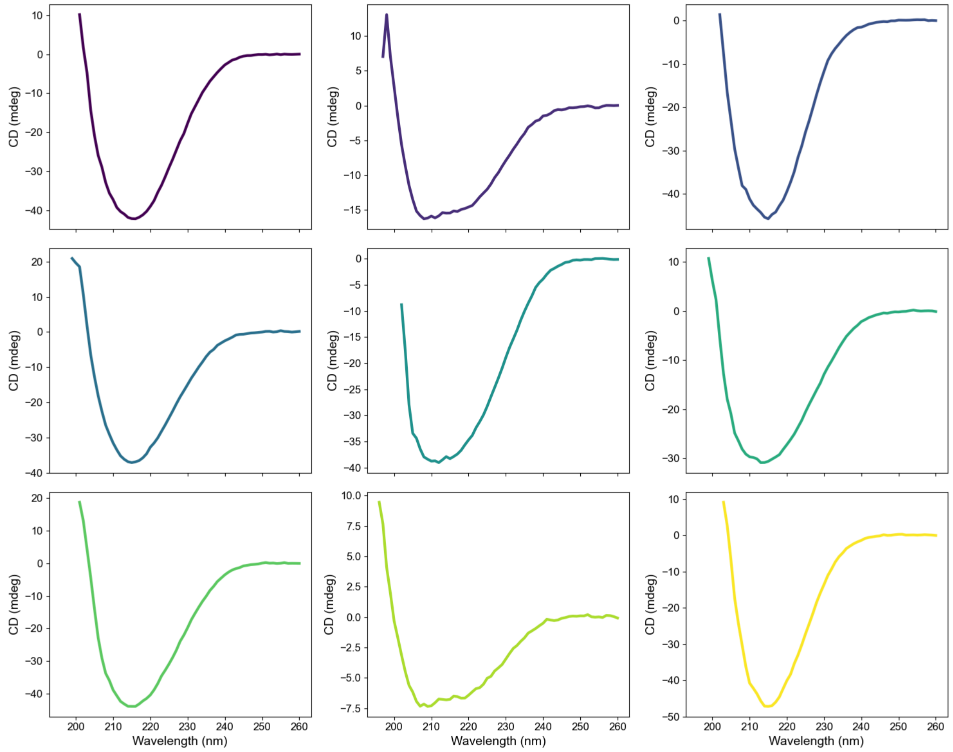


**Figure S27. Circular dichroism (CD) analysis of TMB designs**. CD spectra are shown for the following designs, ordered left to right across each row: TMB_N12_des2, TMB_N16_des4, TMB_N10_des2, TMB_N14_des1, TMB_N12_des3, TMB_N14_des2, TMB_N12_des11, TMB_N16_des5, and TMB_N10_des3.

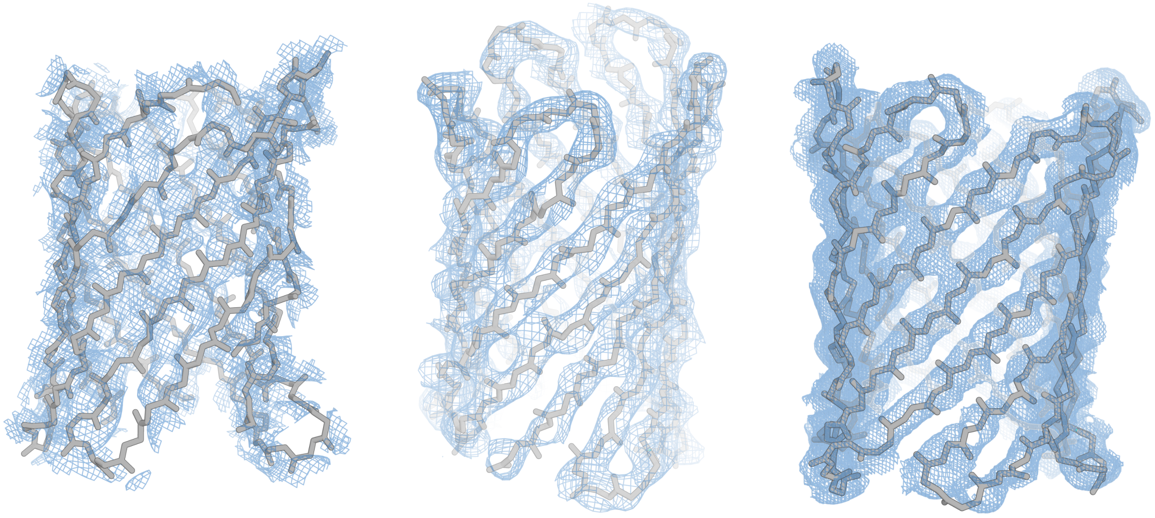

**Figure S28. Electron density maps for crystal structures of TMB designs.** TMB_N10_des2, TMB_N10_des3, and GlyTwistScaffold_des1 are shown from left to right with main chains represented as gray sticks. 2mF_o–DF_c electron density maps contoured at 1.5σ are shown in blue.

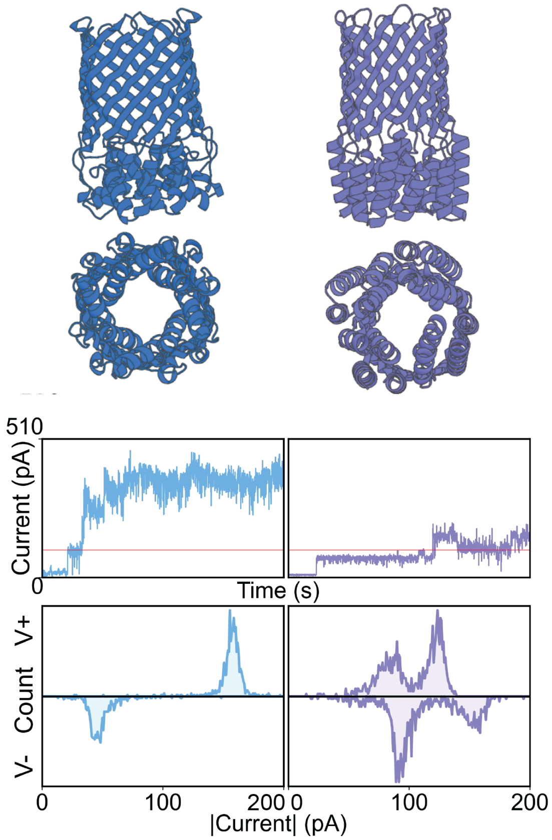


**Figure S29. Structural models and functional characterization of symmetry-enforced TMB designs with extracellular domains.** Top: side and top views of representative symmetry-enforced TMB designs containing extracellular domains. Middle: representative current traces showing multiple pore insertions in 1 M KCl. Bottom: single-channel conductance histograms derived from recordings at ±100 mV

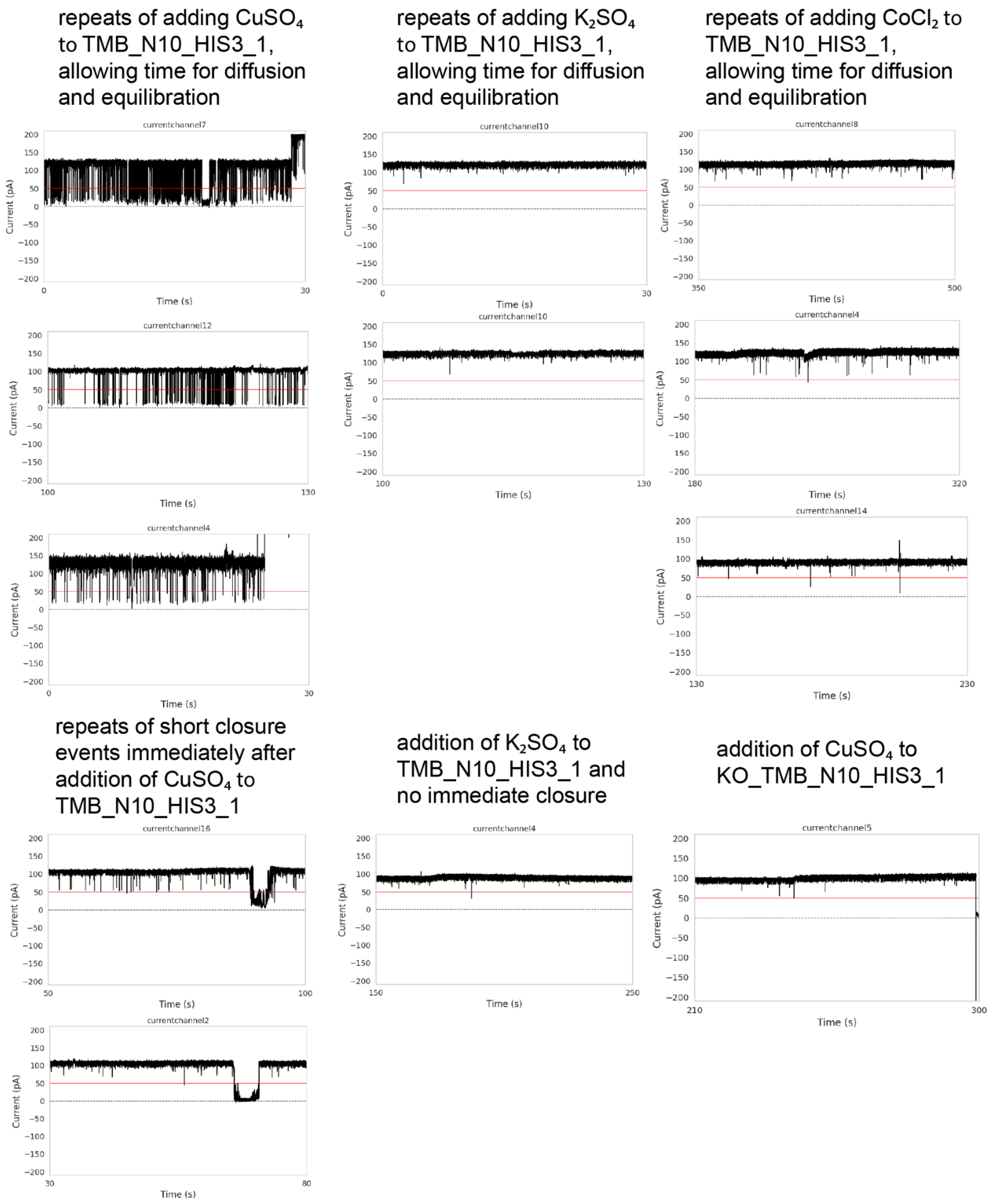


**Figure S30. Reproducibility and control experiments for the histidine-motif design TMB_N10_HIS3_1.** Representative current traces are shown for TMB_N10_HIS3_1 under different experimental conditions, along with a current trace for the histidine-motif knockout KO_TMB_N10_HIS3_1 after addition of $CuSO_4$.

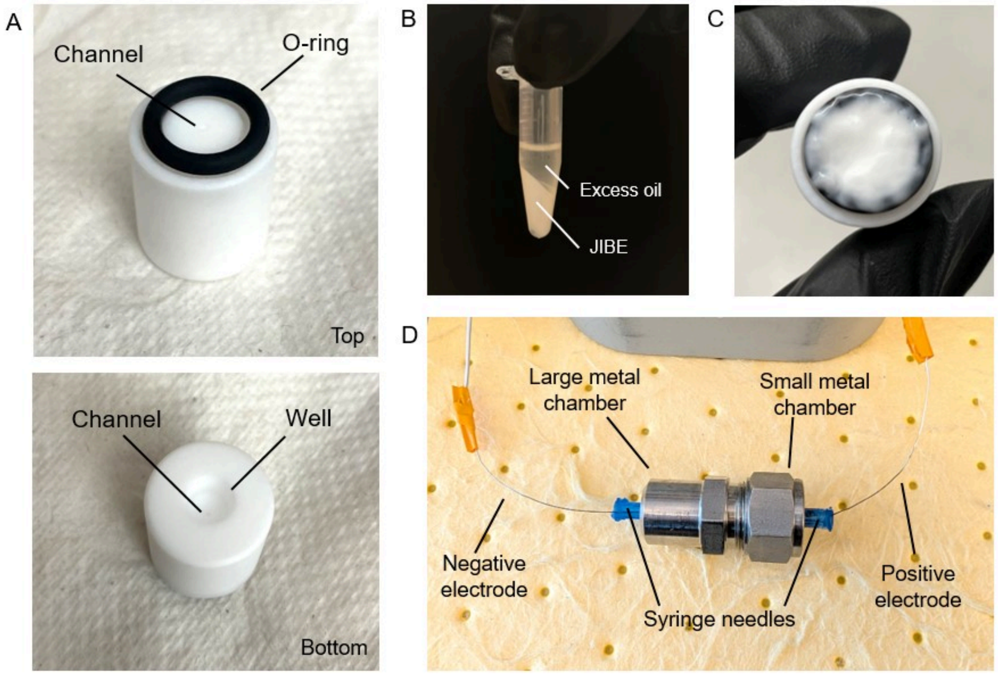


**Figure S31. Overview of the JIBE chamber setup. (A)** Top and bottom views of the plastic chamber with O-ring. **(B)** JIBEs formed with an excess oil layer, which was removed by pipetting before addition to the chamber setup. **(C)** JIBEs covering the surface of the agarose pad and O-ring. **(D)** Fully assembled electrode and chamber setup.

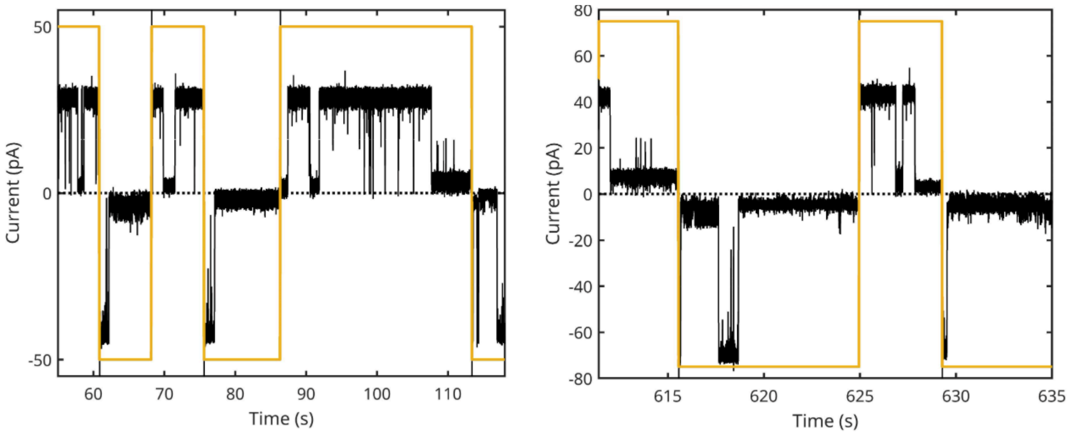


**Figure S32. Stochastic gating of TMB_N14_des13.** Representative current traces show stochastic gating behavior at applied voltages of ±50 mV and ±75 mV (dark yellow lines indicate the applied voltage).
